# Repeatability of adaptation in interacting species

**DOI:** 10.64898/2026.04.01.715933

**Authors:** Loraine Hablützel, Ailene MacPherson, Claudia Bank

## Abstract

In many evolving systems, mutations have background-dependent fitness effects due to genetic interactions between loci within a genome (intragenomic epistasis). In coevolving species, genetic interactions can span across genomes, which has been described as intergenomic epistasis. It is known that intragenomic epistasis can make adaptation more repeatable by constraining accessible mutational paths from a specific starting genotype, generating path repeatability. Here, we investigate how endpoint repeatability, how often adaptation from different starting genotypes reaches the same states in the genotype and fitness space, is influenced by the interplay of intra- and intergenomic epistasis. To this end, we model a two-species community in which the fitness of each depends on the combination of genotypes in both species. We implement this system by constructing and simulating adaptation on coevolutionary fitness landscapes using an *NKC*(*S*) model, an extension of Kauffman’s *NK* model of evolution that includes genetic interactions between loci across species. On these fitness landscapes, we simulate adaptive walks that alternate between mutation in a focal and a partner species. To quantify the repeatability of adaptation, we track the endpoints of adaptive walks and record the fitness of both species at the evolutionary endpoints. We find that intergenomic epistasis causes high endpoint repeatability within a given landscape but highly varying fitness outcomes across different coevolutionary landscapes. Thus, patterns of repeatability deviate from expectations based on single-species adaptation in the presence of intragenomic epistasis due to varying fitness trade-offs between species, which can lead to cycling and large coevolutionary fitness loads.

## Introduction

A longstanding question in evolutionary biology is understanding how repeatable the adaptive processes we observe in nature. Given the same environmental conditions, will populations adapt in the same way? If not, how much of this is due to the stochastic nature of evolutionary processes (for example, mutation or genetic drift). What role do selection and the structure of the genotype space play in determining adaptation? Repeatability is often conceptualised as a measure of the balance between evolutionary potential and constraint. Evolutionary potential can be understood as the capacity to respond to selection, e.g. influenced by the mutation rate, standing genetic diversity, or the fixation probability of beneficial mutations (e.g., Gandon and Michalakis, 2002; Masel and Trotter, 2010). Its counterpart is evolutionary constraint, which can be imposed by pleiotropy, epistasis and the discrete nature of the genotype space that must be traversed in mutational steps (Bank, 2022; Connallon & Hall, 2018; Walsh & Blows, 2009). Genetic interactions (i.e., epistasis) in the discrete genotype space can increase repeatability of evolution by constraining the number of beneficial mutational paths (i.e., path repeatability; Bank, 2022; Weinreich et al., 2006). A different kind of repeatability, which we call endpoint repeatability, measures how frequently the same end states are reached in a finite genotype space. How is endpoint repeatability affected by genetic interactions within and between species, i.e., by intra- and intergenomic epistasis?

By intergenomic epistasis, we refer to systems in which an individual’s fitness depends not only on its own genotype but also on the genotypes of other individuals present in the system, which it interacts with (Hablützel & Bank, 2025; Wade, 2007). Such genetic interactions are intuitive to picture in host-parasite systems, where mutations in virulence and resistance genes can have strong fitness effects on both interacting partners (Carius et al., 2001; Lambrechts et al., 2005; Peever et al., 2000; Salvaudon et al., 2005; Webster et al., 2004). Although researchers have studied the effects of both species interactions (e.g., see Jacquin et al., 2016; Miller et al., 2019, and Venkataram and Kryazhimskiy, 2023 for a review on the repeatability of community-level properties) and intragenomic genetic interactions (e.g., see Bauer and Gokhale, 2015; Li and Zhang, 2025; Szendro, Franke, et al., 2013) on the repeatability of adaptation, few studies have addressed these jointly (but see Burmeister et al., 2016; Gupta et al., 2022). In this study, we use a theoretical approach to explicitly investigate how intergenomic epistasis affects the endpoint repeatability of adaptation.

A common way to study the effects of genetic interactions on adaptation is with fitness landscapes. Fitness landscapes map genotypes to fitness, and the resulting maps give insight into the accessibility of different adaptive paths, as the spread of beneficial mutations moves a population across the fitness landscape towards its peaks (Bank, 2022; de Visser & Krug, 2014; Fragata et al., 2019). Most fitness landscapes are constructed based on genetic interactions within a genome (resulting in a static, single-species fitness landscape). The *NKC* model, developed by Kauffman and Johnsen (1991), captures the effects of both intra- and intergenomic epistasis on fitness by constructing landscapes that depend on the genetic background of another species. In this model, interactions between loci within and between genomes can be tuned separately using different parameters to build randomly generated (i.e., probabilistic) fitness landscapes. The central idea of the *NK* fitness landscape family, to which the *NKC* model belongs, is to capture complex epistatic interactions by randomly drawing fitness components (Kauffman & Johnsen, 1991; Kauffman & Levin, 1987; Kauffman & Weinberger, 1989). Thus, the *NKC* model allows us to implement intra- and intergenomic epistasis without making assumptions about the underlying mechanisms, thereby providing a null model of coevolution.

Fitness landscape models allow us to study repeatability in multiple ways. Here, we assess endpoint repeatability at two levels: (i) the number of realized long-term outcomes of adaptive walks on the same fitness landscape starting from different genotypes, (i.e., stable-state endpoints of adaptation and continuous cycling), and (ii) how frequently these different outcomes are reached (*cf.* Szendro, Franke, et al., 2013). Whereas adaptive walks on single-species landscapes must reach one monomorphic peak genotype (and thus, the number of possible outcomes of adaptive walks is equivalent to the number of peaks of the landscape), multi-species fitness landscapes allow for more dynamic outcomes, including coevolutionary cycling.

Like intragenomic epistasis, we expect intergenomic epistasis to constrain adaptation, e.g. by introducing inter-species contingencies like in Gupta et al. (2022). In their study, Gupta et al. (2022) found that key innovations in *λ*-phage genomes were only achieved if they were preceded by a certain set of mutations in both the *λ* and the *Escherichia coli* genomes. The effect of intergenomic epistasis on re-peatability may depend on the type of biotic interaction. Coevolutionary fitness landscapes dominated by antagonistic interactions may be expected to hinder adaptation, reducing the potential fitness gains of one or both species during adaptation, and increasing the variability in adaptive outcomes if different paths lead to different outcomes (Gupta et al., 2022). Alternatively, fitness trade-offs between species, where some genotype combinations are beneficial for one species but detrimental for another, could promote cycling, helping both species to reach high fitness, but only temporarily (Decaestecker et al., 2007). Finally, landscapes dominated by synergistic interactions may facilitate adaptation and increase endpoint repeatability when mutations in both species increase the fitness of each other.

To test these predictions, we use a minimal version of the *NKC* model with two haploid species, each with two diallelic loci. We generate **coevolutionary fitness landscapes** (see glossary in Table 2) consisting of paired genotypes, where the topography of the landscapes is tunable based on the presence of (i) intergenomic epistasis, (ii) intragenomic epistasis, and (iii) the structure of genetic interactions via the interaction network of loci. We alter the shape of the fitness landscapes via these three parameters to study how the landscape structure affects endpoint repeatability of adaptation (Fig. 3a). We quantify the shape of the fitness landscapes with summary statistics such as landscape **ruggedness**, which captures the non-additivity of fitness effects (Bank et al., 2016; Szendro, Schenk, et al., 2013), and the correlation in fitness ranks between species. Adaptation occurs via adaptive walks of single mutational steps between monomorphic (meta-)genotypes. We alter how species navigate fitness landscapes by varying mutation rates between species and by studying different types of adaptive walks (e.g., random, true and greedy walks). The mutation rate and the type of adaptive walk are both essential parts of a species’ evolutionary potential and determine how quickly a species can find beneficial mutations, by altering their frequency and fixation probabilities, respectively (Fragata et al., 2019; Gandon & Michalakis, 2002).

By allowing the fitness of a species to explicitly depend on the genetic background of the community, we incorporate species interactions into the study of evolutionary repeatability in discrete genotype space. We find that the presence of intergenomic epistasis leads to highly repeatable, yet landscape-specific patterns of adaptation that include cycling between adaptive states and frequently result in fitness costs in one or both interacting species at the evolutionary endpoint. Furthermore, we observe that the interaction structure of the generated landscapes is more important in determining the outcome of adaptation than the manner in which the landscapes are navigated (i.e., the adaptive walk type).

## Model

### Constructing coevolutionary fitness landscapes

To model intra- and intergenomic epistasis, we use the *NKC*(*S*) model by Kauffman and Johnsen (1991). We study a community of two haploid species *X*, where *X* ∈ {F, P}. Throughout the text, we use F to denote the focal species and P for the partner species, following the terminology of Hablützel and Bank (2025), see Table 1 for a full list of notations. Thus, we investigate the smallest possible community for coevolution, where each species has *S* = 1 interacting partners. Each species has *N* loci with *A* alleles at each locus. Throughout, we focus primarily on small genetic networks consisting of *N* = 2 diallelic loci with *A* = 2 for each species unless stated otherwise. We denote the genome of the focal and partner species by the vectors F⃗ and P⃗, respectively. The fitness of a given species *X*, *W* (*X* | ^{⃗^F, P⃗ }) is given by the average of *N* ‘fitness contributions’, one contribution for each locus in its genome (see Fig. 1). The fitness contribution of each locus *i* (*i* ∈ {1*, …N*}) in species *X* is denoted by 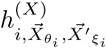 and captures three effects. First, this fitness contribution captures the additive effect of locus *i* on the fitness of species *X*, which depends on the allelic state of this species’ genome at locus *i*, *X⃗_i_*. Second, it captures intragenomic epistasis, which depends on the state of the species’ genome at each of *K* interacting loci, the vector 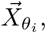 where the set *θ_i_* (|*θ_i_*| = *K*) denotes the set of positions in the species’ genome with which locus *i* interacts. Finally, it captures the effect of intergenomic epistasis, which depends on the allelic state of *C* loci in the genome of the interacting species as denoted by the 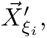 where the set *ξ_i_* (|*ξ_i_*| = *C*) is the set of positions in the interacting genome with which locus *i* interacts. The extent of epistasis is hence determined by two parameters *K* ∈ {0*, …N* − 1} and *C* = {0, 1*, …N*} giving the number of intragenomic and intergenomic interactions, respectively (e.g., *K* ∈ {0, 1} and *C* ∈ {0, 1, 2} for our primary two-locus case). We draw all fitness contributions, 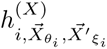 at random from a normal distribution with mean 0 and standard deviation 0.25, N (0, 0.25). Fitness contributions are stored in a look-up table. For each species *X* ∈ {F, P}, there are a total of *A^K^*^+(^*^S×C^*^)+1^ possible fitness contributions for each of the *N* loci in a species’ genome. The resulting look-up table has *N* columns and *A^K^*^+(^*^S×C^*^)+1^ rows, where each row corresponds to a different genetic background representing all possible allele combinations at the relevant loci *X⃗_i_, X⃗_θ_* and 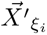 (e.g., Fig. 1). Each fitness contribution can then be retrieved via its unique row ID corresponding to its genetic background (see Supplement A for more details).

**Figure 1:**
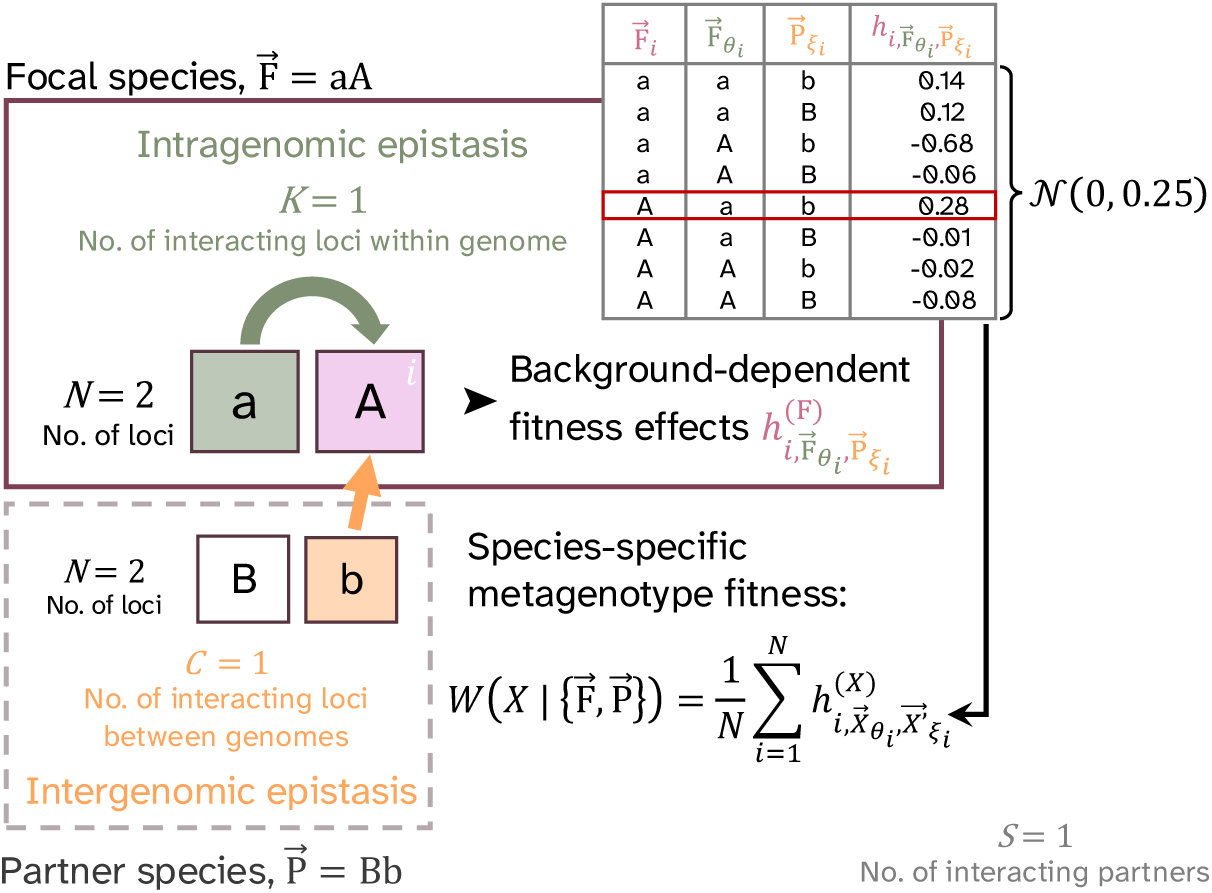
A visual representation of how to compute the fitness contribution of a locus *i* in our model. Each species *X* has *N* = 2 loci with *A* = 2 alleles each. Alleles are encoded as F⃗*_i_* ∈ {a, A} for the focal species and P⃗*_j_* ∈ {b, B} for the partner species. Each locus *i* (here *i* = 2 is highlighted in pink) interacts with *K* loci in its own genome via intragenomic epistasis (green) and *C* loci in the genome of its partner species via intergenomic epistasis (orange). Depending on the combination of alleles at all relevant loci 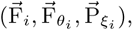 the fitness contribution of locus 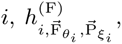, differs. There are *A^K^*^+(^*^S×C^*^)+1^ possible values for 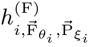 where each value is drawn from a normal distribution and associated with a different genetic background (see top right table). The fitness of a species is then calculated as the average of all *N* fitness contributions in the genome.

**Table 1:**
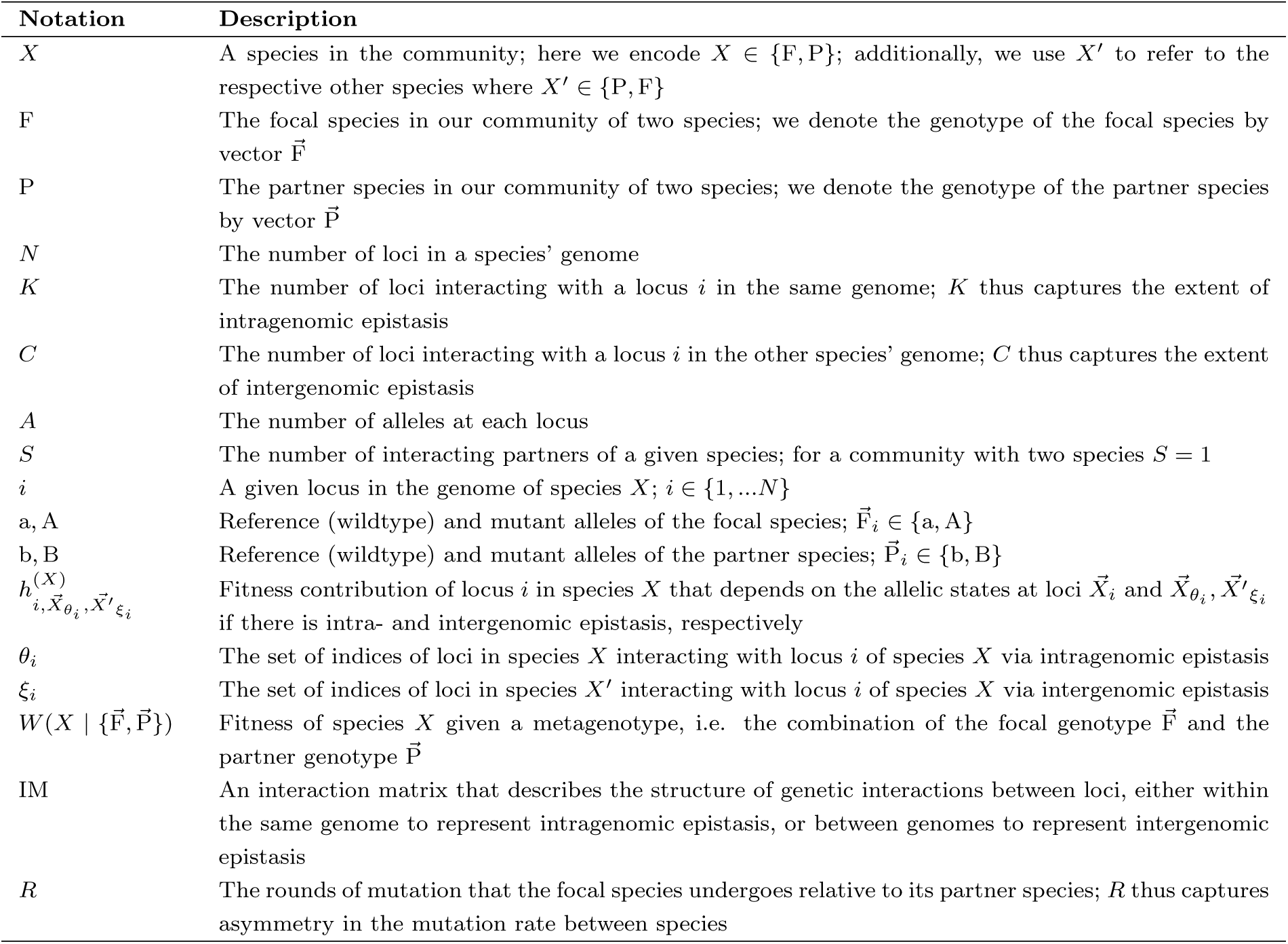
Glossary of the variables and parameters used in the model. Throughout, we refer to the combination of focal and partner genotypes {F⃗, P⃗ } as a “metagenotype”. The number of metagenotypes in a landscape is given by *A*^(*S*+1)^.

**Table 2:** Glossary of fitness landscape terms used throughout the text.

| Term | Definition |
| --- | --- |
| Coevolutionary fitness landscape | The joint fitness landscapes of interacting species coupled through intergenomic epistasis. |
| Fitness rank | The rank of a metagenotype based on fitness across all possible metagenotypes. Fitness rank 1 has the highest (“best”) fitness. |
| (Adaptive) outcome | The state of a community or species at the end of an adaptive walk. Outcomes can either be stable-state endpoints at a (local) fitness peak, or continuous cycling. |
| Fitness load | The reduction of a species’ fitness relative to its hypothetical maximum fitness (i.e., fitness rank -1). In this study, fitness load is caused by intra- and intergenomic epistasis, which prevents species from reaching their global fitness peak. In the latter case, we refer to this as coevolutionary fitness load. |
| Fitness valley | Low fitness (meta)genotypes separating a local fitness peak from higher fitness peaks. |
| Global (fitness) peak | The metagenotype with fitness rank 1, where a species reaches its highest fitness. |
| Joint (fitness) peak | A metagenotype that is a fitness peak for both species. Adaptive walks will stop once they reach a joint peak. |
| Local (fitness) peak | A metagenotype of high but not maximum fitness on a species’ fitness landscape; a local peak is surrounded by single-step mutant neighbours of lower fitness on the species’ landscape but not necessarily on the coevolutionary fitness landscape where fitness is coupled to the partner species. |
| Ruggedness | In this study, ruggedness is understood as the ratio of roughness to slope on a fitness landscape. With increasing ruggedness, the number of peaks on a single-species landscape increases, increasing the likelihood that a species gets stuck on a local peak, rather than reaching the global fitness peak. |
| Sublandscape | The part of a species’ fitness landscape that a species can navigate by itself through mutations in its own genome, which consists of all metagenotypes with the same genetic background in the interacting species. |

The interaction structure, i.e., which loci affect a focal locus *i* (defined by *θ_i_* within a species and *ξ_i_* between species), is determined by interaction matrices IM of dimension *N* × *N*. The rows in these matrices correspond to the *N* loci of a species *X* and the columns either correspond to the same *N* loci of *X* for intragenomic interactions, or the *N* loci in the other species *X^′^* for intergenomic interactions. The entries of these matrices encode interactions by 0s and 1s, where 1 represents an interaction and 0 represents no interaction. For example, the intragenomic interaction matrix in the focal species with *N* = 2 and *K* = 0 reads as

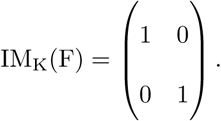

Here, all off-diagonal elements are 0 because *K* = 0 indicates the absence of intragenomic interactions, whereas the 1 in the diagonal indicates the additive fitness effects. Interspecific interactions are characterised by pairs of such matrices, one capturing the interaction of each locus in the focal species with loci in the partner species, and the other capturing the interactions of each locus in the partner species with loci in the focal species. We study the effect of intergenomic interaction structure by considering all four different categories of such matrices (see Supplement A for more details).

Finally, we combine the look-up tables and interaction matrices to construct the fitness landscapes for each species *X* ∈ {F, P} in the community. For this, we compute the fitness of all *A^N^*^+^*^N^* possible metagenotypes – the genotype combinations of the focal and the partner species as denoted by {⃗F, P⃗ }. Each metagenotype {⃗F, P⃗ } maps to two fitness values that indicate the fitness of the focal and partner species, respectively, given by:

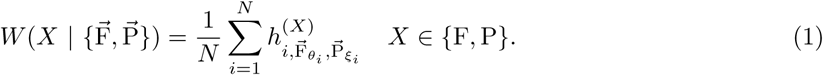

This model thus generates a two-dimensional fitness space for a system of two species (i.e., mapping *A^N^*^+^*^N^* → ℝ^2^; or *A*^(^*^S^*^+1)*×*^*^N^* → ℝ^(^*^S^*^+1)^ for a system with *S* interacting partners per species). Fig. 2 shows an example of a two-dimensional fitness mapping generated for a pair of interacting species with *N* = 2 loci and *A* = 2 alleles. For each parameter combination of *K* and *C*, we generated 200 fitness landscapes. For *C* = 1, we used four different intergenomic interaction matrices and generated 50 fitness landscapes for each type. To compare evolutionary outcomes across landscapes, we recorded the **fitness ranks** of adaptive outcomes, rather than the fitness values of these outcomes. Due to the random nature of the fitness contributions in our models the range in fitness values among genotypes can vary substantially across landscape replicates and hence can not be compared directly.

**Figure 2:**
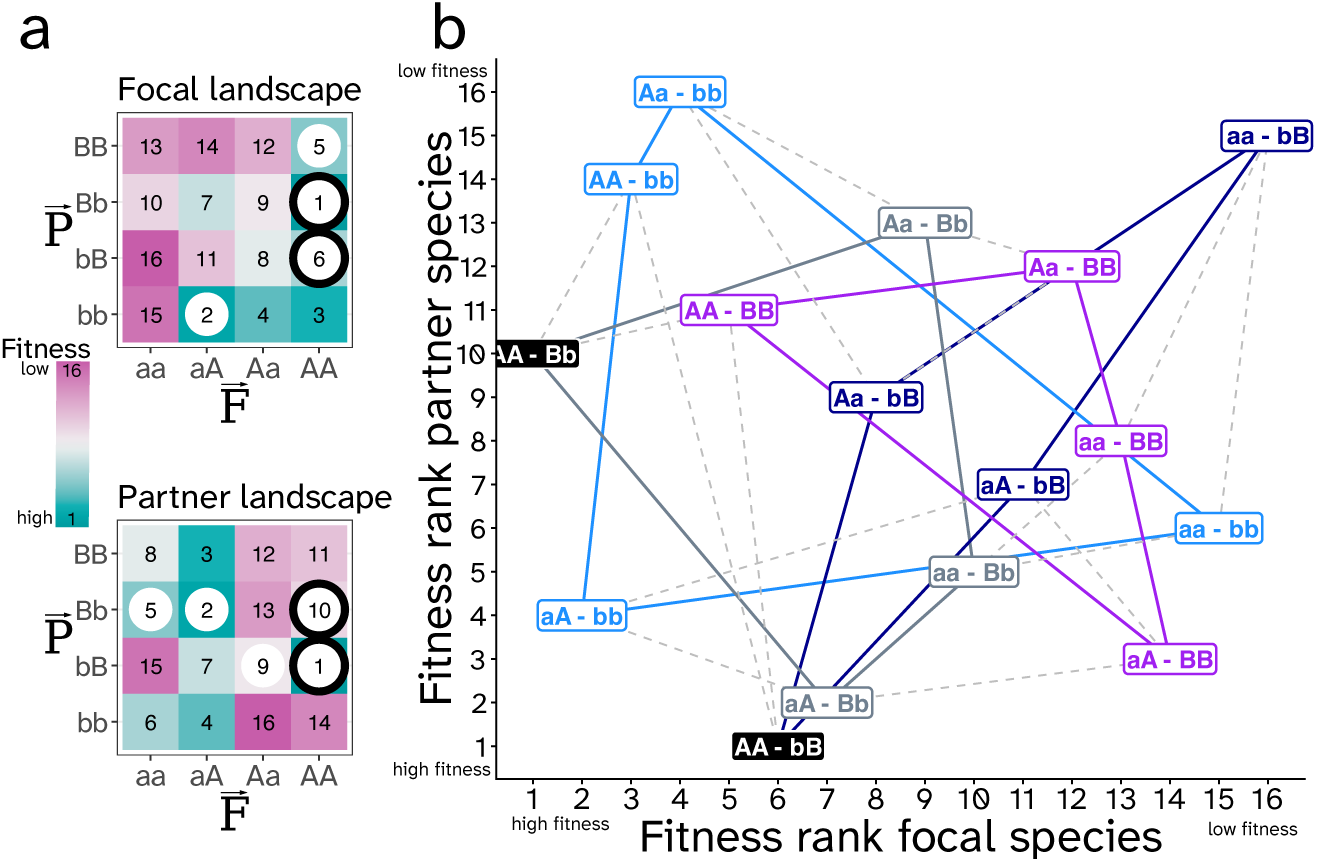
Two ways of visualising a coevolutionary fitness landscape with *N* = 2*, K* = 1*, C* = 1*, A* = 2*, S* = 1. Alleles are encoded as F⃗*_i_* ∈ {a, A} for the focal species and P⃗*_i_* ∈ {b, B} for the partner species. Here, each genotype combination (i.e., each metagenotype {F⃗, P⃗ }) confers a unique fitness value for each species and is ranked from 1 (global fitness peak) to 16. In **a)**, the coevolutionary landscape for both species viewed separately where local species-specific fitness peaks are indicated with white circles. Local species-specific fitness peaks are genomic combinations where both possible mutations in the species of interest reduce fitness. Metagenotypes with a black outline fulfil the peak condition for both species and are thus **joint fitness peaks**. The landscape shown has two joint fitness peaks caused by reciprocal sign epistasis in the partner genome on the AA genetic background. **b)** the two-dimensional fitness mapping of all metagenotypes. Colored solid lines show the different **sublandscapes** of the focal species that can be navigated via mutations in the focal genome while keeping the partner species’ genome unchanged. Dashed lines show mutations in the partner genome that can move the focal species to a different sublandscape as the intergenomic background changes. We highlight the two joint fitness peaks in black.

**Figure 3:**
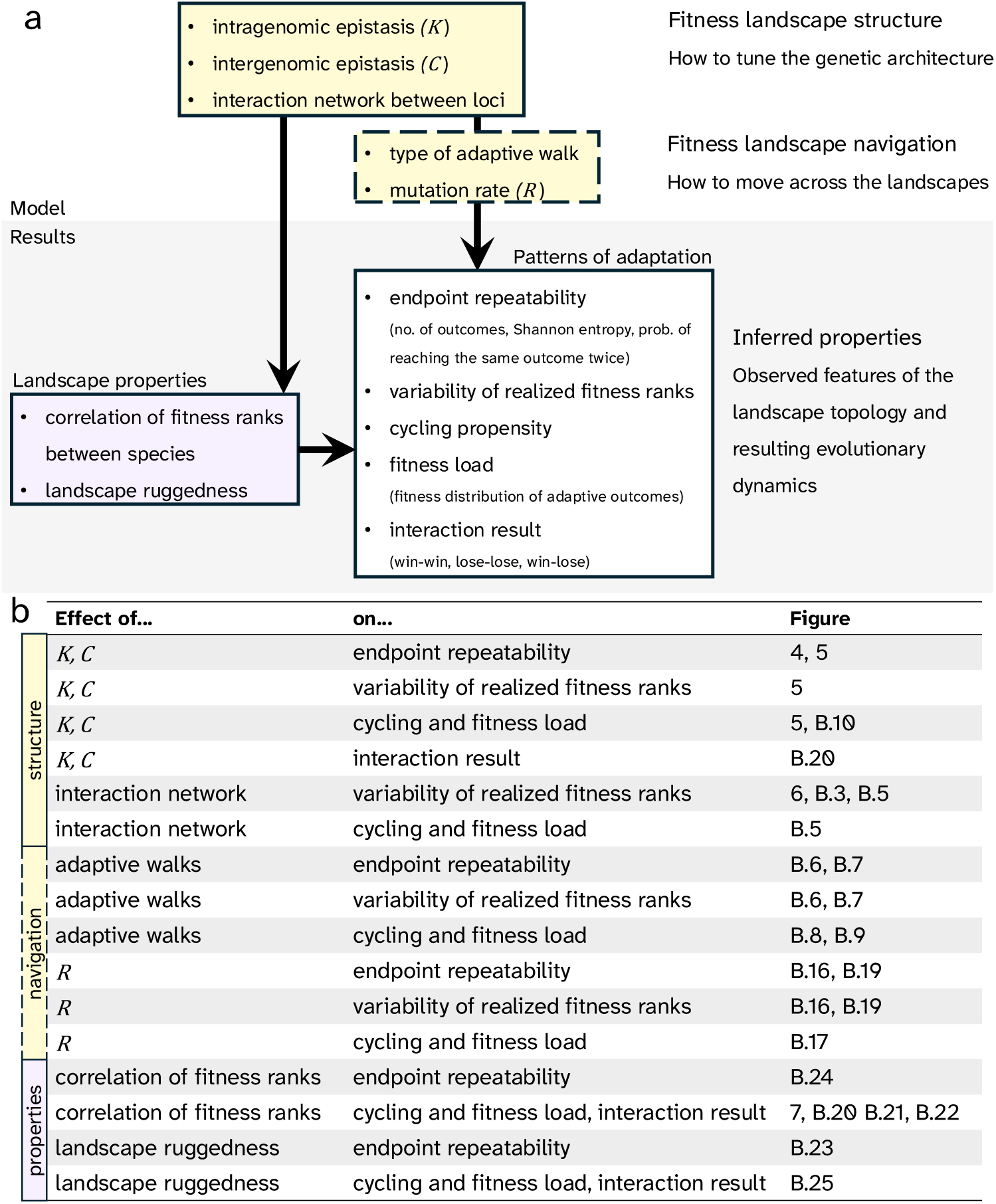
a) A visual overview of the model with the main variables we investigate (yellow boxes) and the landscape properties we infer and measure (underlaid in grey). On the one hand, we manipulate intra- and intergenomic epistasis and thus the shape of the fitness landscapes, and on the other hand, we vary how species navigate across the landscapes during adaptive walks. We then assess how the shape and navigation of the fitness landscape impact patterns of adaptation, such as endpoint repeatability. Additionally, we infer several landscape properties and study how they relate to the observed patterns of adaptation (purple box). In **b)**, we list the main relationships investigated in the manuscript and in which figures the results are shown.

### Comparison with non-coevolutionary fitness landscapes

To provide “abiotic” equivalents to the dynamic *NKC*(*S*) fitness landscapes with *S* = 1 interaction part-ners and *N* = 2 loci, we implemented two single-species fitness landscape models. Firstly, we used an *NK* model with *N^′^* = *N* + *N* loci and *K^′^* = *K* + *C* genetic interactions between loci, which otherwise follows the same model laid out above. Since all genetic interactions in the *NK* model are intragenomic, this helps contrast intergenomic epistasis to classical expectations based on intragenomic epistasis. However, the fitness dimensionality of this model is different from the *NKC*(*S*) fitness landscape: *A^N^*^+^*^N^* → ℝ.

Secondly, we introduced a single-species seascapes model, which we denote as the *NK*(*E*) model, where the fitness values of a species’ genotypes change in a time-dependent manner when switching between different abiotic environments (inspired by Merrell, 1994; Mustonen and Lässig, 2009). Again, we used *N^′^* = *N* + *N* loci, but in this model, the genome is divided into *E* blocks of *N* loci that each encode a separate trait under selection in one of two environments (*E* = 2). Thus, in the *NK*(*E*) model, the focal species’ fitness *W* (F | F⃗*, Y*) depends on the genotype F⃗ and the environment *Y*. Effectively, instead of modelling coevolution between two different species, this seascape implementation models coevolution between two traits that are under selection in two different environments. This maintains the dimensionality of the fitness mapping in the *NKC*(*S*) landscapes, because the same genotype can differ in fitness between environments: *A^N^*^+^*^N^* → ℝ^2^. Furthermore, the parameters *K* and *C* of the *NKC*(*S*) translate into genetic interactions between loci within the same trait (*K_w_* ≡ *K*) or between different traits (*K_b_* ≡ *C*) in the *NK*(*E*) model. See Supplementary Section A for more information on the *NK*(*E*) model and comparisons between the three models.

### Adaptive walks

To simulate adaptive walks on the coevolutionary fitness landscapes with a focus on endpoint repeatability, we initiated our two-species system at each of the metagenotypes on the landscapes. Both species were then given alternating turns to navigate their landscape using fitness-increasing single-mutation steps. Adopting a strong selection, weak mutation (SSWM) regime, we assumed that, given that at least one fitter single-step mutational neighbour exists, the whole population instantly moves to the newly chosen metagenotype ^{⃗^F, P⃗ } (e.g., moving along the lines in the direction of increasing fitness rank in Fig. 2). In the main text, we show the results for random adaptive walks, in which the probability of fixing a fitness-increasing single-step mutation is the same for all fitness-increasing single-step mutations accessible from the current genotype (but see Supplement B for results for true and greedy adaptive walks and Fragata et al. (2019) or Supplement A for detailed explanations of the respective algorithms). As the focal species mutates its genotype on the fitness landscape, increasing its fitness *W* (F | {⃗F, P⃗ }) through accessible mutational steps (i.e., mutating one of its own *N* loci and moving along the solid lines in Fig. 2b), this alters the metagenotype {⃗F, P⃗ } of the community and thus updates the fitness *W* (P | {⃗F, P⃗ }) of the partner species. This can either increase or decrease the partner species’ fitness *W* (P | {⃗F, P⃗ }) if *C* ≠ 0. Mutations in the focal species have no effect on *W* (P | {⃗F, P⃗ }) if *C* = 0, because transitions between metagenotypes are neutral when there are no intergenomic interactions. Next, the partner species is allowed to mutate (e.g., moving along the dashed lines in Fig. 2b). An adaptive walk is a series of alternating steps on the fitness landscape and terminates once both species reach a fitness peak or when the walk reaches the maximum number of steps (arbitrarily set to 300). The **outcome** of an adaptive walk can either be categorised as stable-state endpoint of adaptation (a joint fitness peak), or as cycling behavior when walks exceed the maximum number of steps. We repeatedly simulated adaptive walks starting from all possible metagenotypes {⃗F, P⃗ }, with each starting metagenotype repeated 5 times.

To explore further variations in fitness landscape navigation, we consider mutational asymmetry between species during adaptive walks. Following Bull (2021), we introduce *R* as the rounds of mutation that the focal species undergoes relative to its partner species. Thus, *R* mimics different mutation rates between the two species by allowing the focal species to take multiple steps on the fitness landscape before the partner species can take a step. We interpret *R* as the mutation rate rather than generation time, due to the non-trivial effects generation time can have on adaptation (Hablützel et al., 2025).

### Implementation

Fitness landscape construction, simulation of adaptive walks, subsequent analysis and visualisation were all performed in *R* (version 4.5.1, R Core Team, 2026). Unless noted otherwise, for each model (*NKC*(*S*), *NK*, *NK*(*E*)) and parameter combination, we assessed repeatability across 200 randomly generated land-or seascapes, including landscapes with different interaction matrices. For each land- or seascape, we simulated 80 adaptive walks, conditioned on the focal species mutating first (5 replicates starting at each of 16 initial (meta)genotypes).

## Results

### The effects of intergenomic epistasis on the repeatability and variability of adaptation

#### Intergenomic epistasis maintains high endpoint repeatability

Comparing adaptation on fitness land- and seascapes with different kinds of genetic interactions (i.e. different combinations of *K*, *C*, *K_w_*, and *K_b_*), we find that the number of adaptive outcomes per landscape replicate remains low with increasing intergenomic epistasis on *NKC*(*S*) landscapes, but grows with increasing intragenomic epistasis (*K*, *K_w_*, *K_b_*; Fig. 4a-c). (Note that in the *NK* model, the number of adaptive outcomes is equivalent to the number of peaks of the landscape.) On completely additive landscapes (*K* = 0, *C* = 0), the smooth nature of the landscape generates a single adaptive outcome, which is the **global fitness peak** that maximises a species’ fitness. With increasing frequency of genetic interactions between loci, non-additive fitness effects generate landscape ruggedness, which is expected to be associated with the appearance of **local fitness peaks** and **valleys**. Adaptive walks on rugged fitness landscapes can stall on these local peaks, which generates alternative adaptive outcomes on a landscape. Although both inter- and intragenomic epistasis increased landscape ruggedness (measured as the ratio of roughness to slope; see Fig. B.1), *NKC*(*S*) landscapes showed fewer adaptive outcomes per landscape compared to their *NK* and *NK*(*E*) counterparts with equivalent ruggedness.

**Figure 4:**
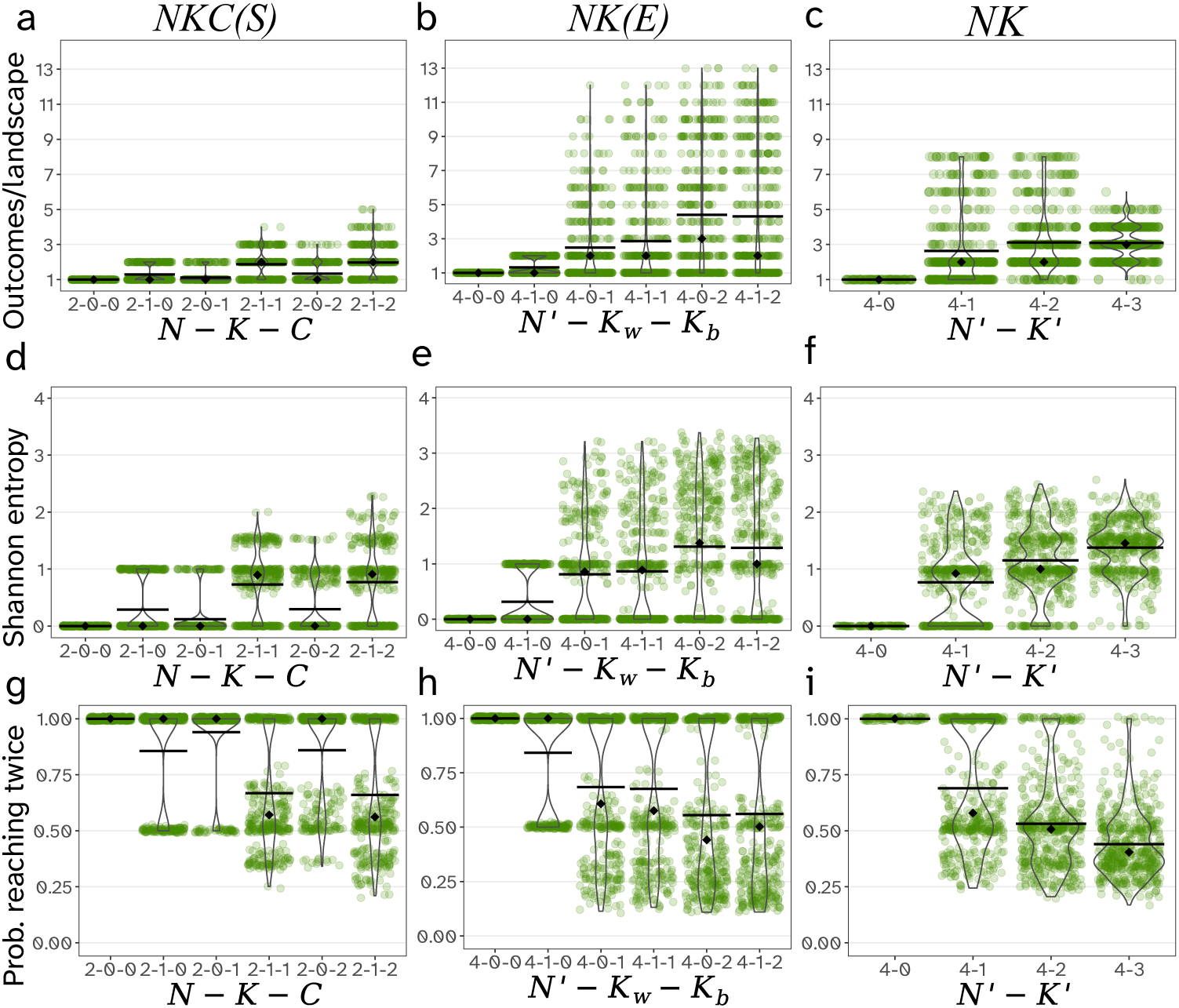
Multiple repeatability proxies indicate that high endpoint repeatability is retained on *NKC*(*S*) landscapes in the face of intergenomic epistasis. For differing combinations of inter- and intragenomic epistasis parameters, we assessed endpoint repeatability for 80 replicated adaptive walks of 200 randomly generated land-and seascapes. Measures of repeatability include the number of observed genetic endpoints (**a-c**), Shannon entropy of adaptive outcomes (**d-f**), and the probability of reaching the same outcome twice on a given landscape (**g-i**). Overall, *NKC*(*S*) landscapes (**a,d,g**) with intergenomic epistasis showed higher endpoint repeatability relative to their *NK*(*E*) (**b,e,h**) and *NK* counterparts (**c,f,i**). Black bars show means, black diamonds show medians.

We used three measures of endpoint repeatability to capture both the number of possible outcomes and their relative frequencies of occurring. We measured (i) Shannon entropy to capture the probability distribution of all possible adaptive outcomes, (ii) the probability of reaching the same outcome twice (following Amado and Bank, 2024), and (iii) the frequency of the most common outcome on a given land-scape. All of these metrics displayed comparable patterns of higher endpoint repeatability on *NKC*(*S*) landscapes as compared with *NK* and *NK*(*E*) landscapes; low entropy of adaptive outcomes (Fig. 4d-f) typically corresponded to a high probability of observing the same adaptive outcome twice on the same landscape (Fig. 4g-i), and the most common outcomes occurred at high frequency (Fig. 5). All measures pointed to high endpoint repeatability in the presence of intergenomic epistasis.

**Figure 5:**
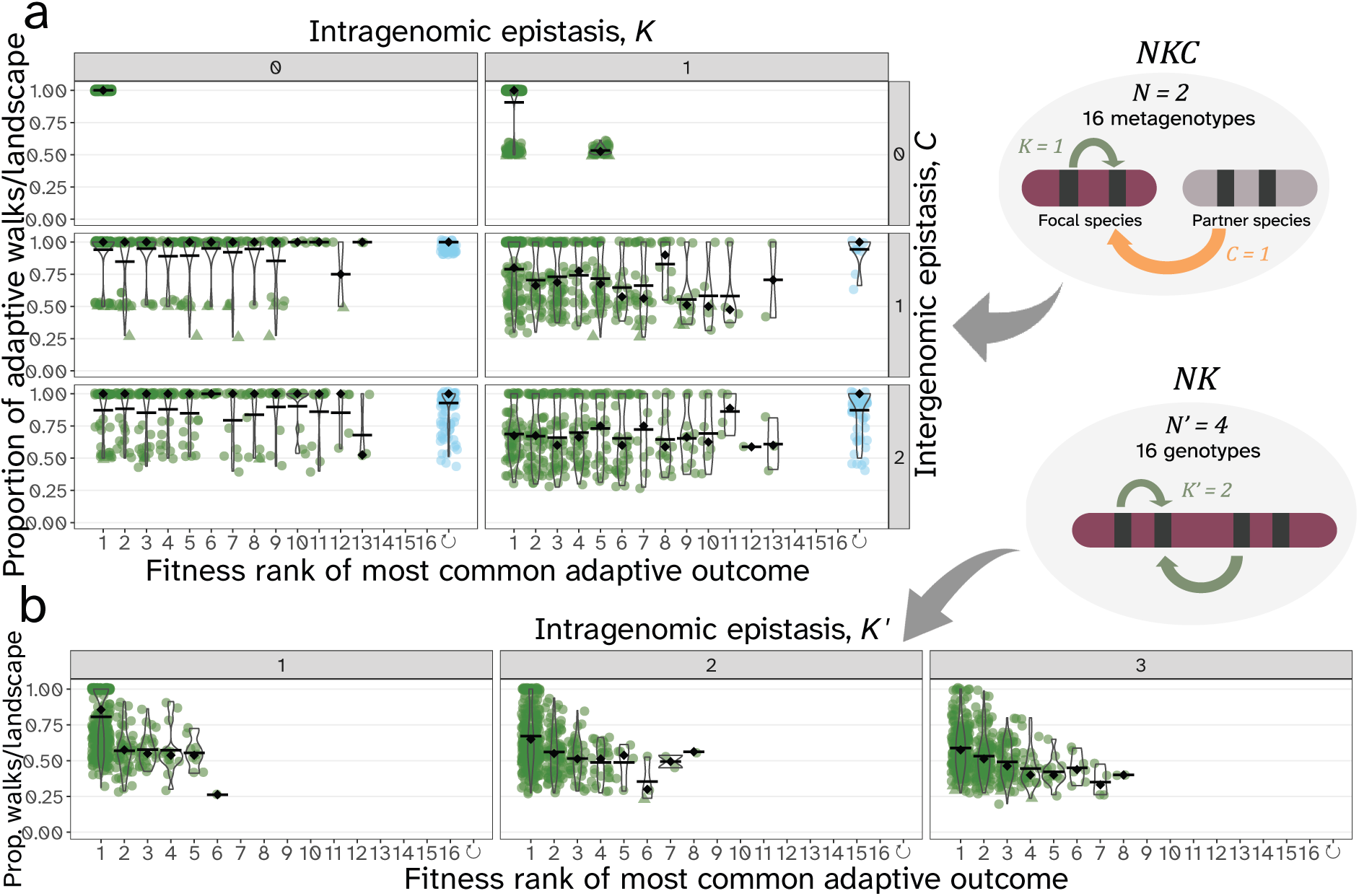
Intergenomic epistasis increases the variation in the realized fitness rank of the most common adaptive outcome across landscape replicates. For analogous *NKC*(*S*) (panel **a**) and *NK* models (panel **c**), we show how often the most common adaptive outcome is reached as a function of its fitness rank and the proportion of adaptive walks that led there. Each landscape replicate is represented twice, once for the focal species and once for the partner species (for outcomes that were tied as most common, we randomly chose one and represent it as a triangle). Black bars show mean, black diamonds show median proportions for each fitness rank. Note that in the *NKC*(*S*) model for *C* = 0, neutral transitions between metagenotypes cause plateaus where several metagenotypes map to the same fitness rank, whereas intergenomic epistasis (*C >* 0) generally introduced unique fitness ranks for all 16 metagenotypes. Outcomes with intergenomic epistasis (*C >* 0) include the possibility of cycling (blue dots). For single-species fitness landscapes with *N^′^* = 4 (panel **b**), frequencies of the most common adaptive outcomes correlated with their fitness ranks across landscape replicates.

#### Intergenomic epistasis causes large variability in adaptation across landscape replicates

In contrast to the observed high endpoint repeatability on a given fitness landscape, *NKC*(*S*) landscapes with intergenomic epistasis (*C >* 0) yielded endpoints with highly variable fitness ranks across fitness landscape replicates (see Fig. 5a). This is because intergenomic epistasis couples the fate of the two interacting species, which reduces the number of (joint) adaptive outcomes while decreasing the possibility that either species can adapt to its global fitness peak. In both dynamic landscape models (*NKC*(*S*) and *NK*(*E*)), the multidimensional fitness mapping imposed additional conditions on which (meta)genotypes could be realized as adaptive outcomes (e.g., see Fig. 2 for an *NKC*(*S*) example). Adaptive walks could only terminate on joint fitness peaks, where a given (meta)genotype was a fitness peak for both interacting species or traits simultaneously. If such joint fitness peaks were not present on a given landscape or inaccessible from certain starting (meta)genotypes, this resulted in continual cycling between multiple (meta)genotypes; an additional long-term outcome of adaptation (blue dots in Fig. 5a and B.12). However, the variability in the realized fitness ranks (and thus, the potential fitness cost) of stable-state adaptive outcomes was more pronounced on *NKC*(*S*) landscapes than *NK*(*E*) seascapes (compare Fig. 5a and B.12, or Fig. B.10a and B.13).

For *NK* landscapes, variability in realized fitness ranks across landscape replicates was low. As expected, intragenomic epistasis introduced local peaks and valleys on the fitness landscapes (i.e., ruggedness), at which adaptive walks could stall instead of progressing to the global fitness peak. However, this pattern was comparable across different fitness landscape replicates, indicating a correlation between the fitness rank of an endpoint and its frequency on a given landscape (Fig. 5b). Thus, the patterns of adaptation was similar across *NK* landscape replicates with the same parameters, whereas different *NKC*(*S*) landscape replicates tended to produce unique patterns of adaptation.

#### Variation in realized fitness ranks depends on the underlying structure of intergenomic interactions

The observed patterns of variability in realized fitness ranks across landscape replicates depended not only on the presence or absence of intergenomic epistasis but also on the structure of the intergenomic interactions. We illustrate this for *C* = 1 in Fig. 6, for four possible interaction networks. We classified these interactions as “congruent” if the interaction matrix of both species is structured the same and “specific” if only one of two loci was involved in the interaction.

**Figure 6:**
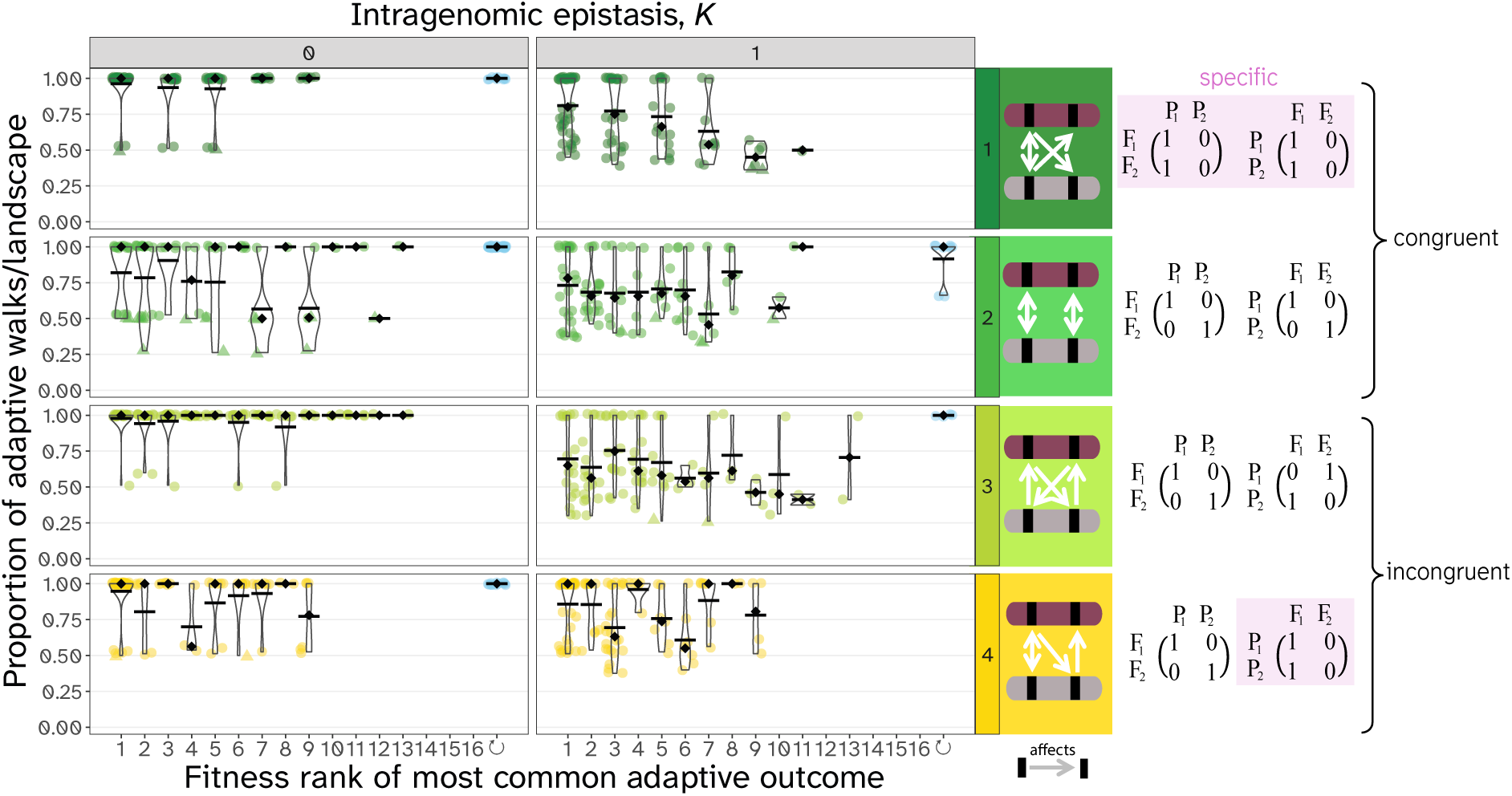
Different interaction structures of intergenomic epistasis affect observed patterns of repeatability and variability. Here, we show the results from Fig. 5a with *C* = 1 seperately for the different underlying intergenomic interaction matrices. For outcomes that were tied as most common, we randomly chose one and represented it as a triangle. Interaction structures and the respective interaction matrices are shown in the panels on the right, where arrows indicate which loci interact with each other (only showing intergenomic epistasis). The investigated interaction networks differ in their congruence and the specificity of interaction between loci. Interactions are congruent if the interaction matrices are identical for both species, and they are specific if they only depend on a single locus in the genome of the other species. Specific interactions introduce neutral plateaus on the coevolutionary landscape where multiple metagenotypes map to the same fitness rank.

Landscapes generated using network 1 (specific and congruent interactions) showed fewer realized outcomes of adaptation across landscape replicates than other interaction matrices. This is because one of the loci in each species is effectively neutral in this interaction, which introduces redundancy between different metagenotypes (i.e. some of the metagenotypes map to the same fitness; Fig. B.2). This pattern holds more generally; specific interactions decreased variability in realized fitness ranks between different fitness landscape replicates by limiting the number of possible fitness outcomes (Fig. B.3). Secondly, we observed more multi-peaked coevolutionary landscapes with interaction networks 2 and 4 than with networks 1 and 3 in the absence of intragenomic epistasis (26% and 16% of landscapes vs. 8% and 6% of landscapes). Interestingly, although different interaction networks produced specific patterns, they did not differ in landscape features such as their ruggedness (Fig. B.4). Different types of interaction structures also affect endpoint repeatability and variability in realized fitness ranks in the *NK* model (Fig. B.15).

#### Repeatability patterns are largely robust to different adaptive walk types

Finally, we examined the robustness of repeatability to differences in how the landscapes are navigated. We found little difference in the observed patterns of endpoint repeatability across different types of adaptive walks (random, greedy, true; see Fig. B.6 and B.7). We then altered the relative mutation rate between the two species by allowing (i) asymmetry and (ii) randomness in the number of subsequent adaptive steps in each species, thereby mimicking different mutation rates for the interacting species, as determined by the parameter *R*. However, neither asymmetric nor random mutation sequences altered the patterns of repeatability or variability in realized fitness ranks (Fig. B.16, B.17, B.19).

### The effects of intergenomic epistasis on the fitness load of adaptive outcomes

#### Intergenomic epistasis can come at a cost of a species’ fitness

Although the endpoints of coevolution were often highly repeatable, Fig. 5a showed that many of these repeated adaptive outcomes were unfavourable for one or both of the interacting species. The global fitness peak was still the most common outcome of adaptation when there was intergenomic epistasis (*C >* 0), but it became less common to reach fitness rank 1 on landscapes with intergenomic epistasis than on comparable landscapes without intergenomic epistasis for *N* = 2 and *N^′^* = 4 (compare Fig. B.10 and B.14). This indicated that there was a **fitness load** associated with intergenomic epistasis that reduced the achievable fitness of coevolving species and communities (Fig. B.11). We defined the fitness load of a species as the difference in fitness rank between each observed adaptive outcome and the global fitness peak, essentially shifting the fitness rank values by 1 for reinterpretation. Consistent with their effect on variability of realized fitness ranks across landscapes, specific interaction networks on average reduced the difference between the distribution of fitness loads between the focal and partner species. Moreover, by adapting on landscapes with a reduced number of possible adaptive outcomes, species with a specific interaction had a higher probability to reach their global fitness peak (both species in network 1; partner species in network 4; Fig. B.5).

#### Inferred species interactions affect winner-loser dynamics of interacting species

The presence of fitness load begs the question of whether both interacting species tend to suffer from the same amount of load or whether there are “winners” and “losers” of coevolution. Though the *NKC*(*S*) model does not allow us to define explicit ecological interactions, we inferred a statistical property of the landscapes as a simple proxy for species interactions. We calculated the rank correlation of the metagenotypes within a coevolutionary fitness landscape, which determines whether metagenotypes that conferred high fitness to the focal species on average also conferred high fitness to the partner species and vice versa. Along this continuum of inferred interactions we classified four different kinds of interaction types: (i) win-win interactions, where both species reached their global fitness peak, (ii) win-lose interactions, where one species reached its global fitness peak, but the other species was at a lower-than-possible fitness peak, (iii) lose-lose interactions, where neither species reached its global fitness peak, and (iv) cycling (Fig. 7a).

**Figure 7:**
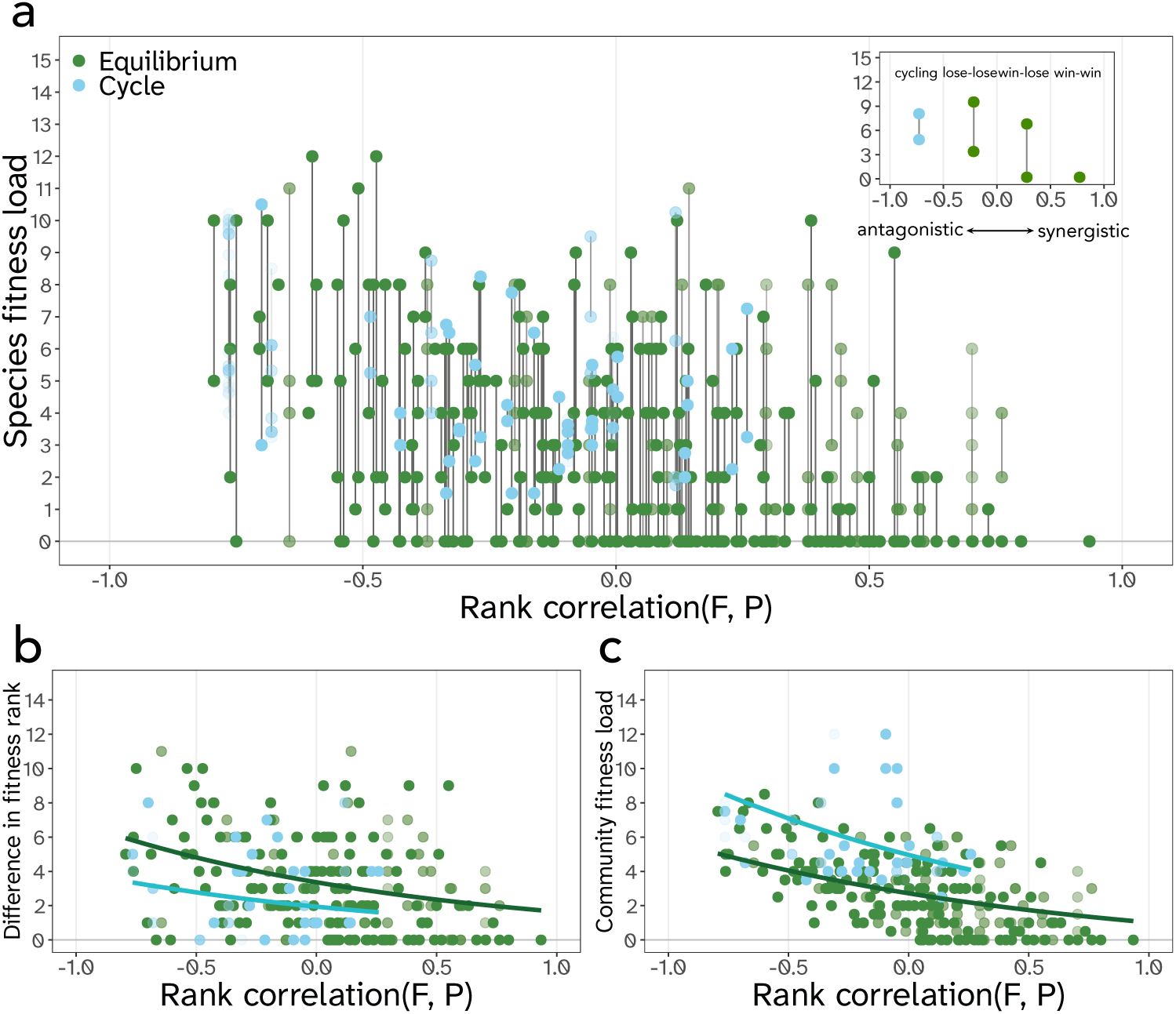
The fitness load of species and communities decreases with the inferred correlation of fitness ranks between species, here shown for landscapes with *N* = 2*, K* = 0 and *C* = 1. In panel **a)**, each dot shows the fitness load of a species at the end of an adaptive walk. Species pairs from the same fitness landscape are connected by a vertical grey line. The opacity of the dots corresponds to their level of endpoint repeatability (i.e., faint dots only occurred for a subset of adaptive walks on a given fitness landscape). Inset: A scheme of interaction results and our a priori expectations of where in the spectrum of inferred interactions they may occur. In panel **b)**, the difference in fitness ranks between species at the endpoint of adaptation (length of grey bars in **a**) decreases only slightly across the ecological interaction continuum as a measure of asymmetry in fitness load, whereas the mean fitness rank difference for cycles remains almost constant across the continuum of inferred rank correlations. Panel **c)** shows community fitness load as defined by the mean fitness load of both species. Cycling results are shown in blue based on the mean fitness ranks of species during the cycle. For **b)** and **c)**, we fitted a generalised linear mixed model and plotted the predicted values as trend lines for comparison of cycles and equilibrium outcomes; see Table B.1 for model results. See Fig. B.20, B.21 and B.22 for respective results on landscapes with other values of *K* and *C*.

We found that asymmetries in fitness load were prevalent and that the different interaction types occurred along much of the rank correlation continuum (Fig. 7a, B.20). However, the extent of the asymmetry in fitness load increased slightly as the correlation between fitness ranks became more negative (Fig. 7b, Fig. B.21). This was also reflected in the trend of the mean community fitness rank, which increased as the rank correlation between species became more positive (Fig. 7c, Fig. B.22). Notably, differences in mean fitness ranks between species tended to be lower in cycles, whereas community fitness load was increased (e.g., Fig. 7b and c). This implies that cycles were overall detrimental for the community, although they buffered the asymmetry in fitness load between the species.

Introducing asymmetry in the mutation rate (*R*) did not alter winner-loser dynamics, against our expectation that an increased mutation rate would increase a species’ probability of navigating the fitness landscape towards its global fitness peak, thus favouring the species with a high mutation rate as a winner of coevolution (Fig. B.18).

#### Inferred landscape features do not predict the repeatability of adaptive outcomes

Finally, we investigated the relationship between several inferred properties of the simulated fitness landscapes and the repeatability of different adaptive outcomes. We found that neither landscape ruggedness (Fig. B.23) nor the rank correlation between species (Fig. B.24) explained variation in endpoint repeatability measured as its frequency on a given fitness landscape. Thus, an inferred overall positive interaction could lead to the same level of repeatability as an inferred negative interaction.

## Discussion

In this study, we used a classic fitness landscape model of coevolution to investigate the effects of intra-and intergenomic epistasis on the repeatability of adaptation in a system of two interacting species. We characterised adaptation on the fitness landscapes from every possible starting point using three main measures: (i) the number of adaptive outcomes per landscape, (ii) their Shannon entropy, and (iii) the relative frequency of the most common outcomes. Moreover, we quantified the fitness load for individual species and the community that is associated with the coevolutionary process. We found that, within a given landscape, intergenomic epistasis led to highly repeated, landscape-specific endpoints of adaptation, often associated with a fitness load for one or both interacting species. Conversely, intragenomic epistasis led to less endpoint repeatability but more consistent results across replicates. Patterns in the repeatability, variation in realized fitness ranks, and the associated fitness load primarily depended on the underlying fitness landscape parameters, i.e. the extent and presence of intra- or intergenomic epistasis (*K* and *C*) and the network of intergenomic genetic interactions.

In both single-species and coevolutionary landscapes, intragenomic epistasis introduces local peaks and valleys on the fitness landscapes of the individual species. In our study, intragenomic epistasis increased the number of realized adaptive outcomes on a given fitness landscape, thereby decreasing endpoint repeatability for individual landscape replicates. Intragenomic epistasis has previously been proposed to increase path repeatability (Bank, 2022; Weinreich et al., 2006), which refers to the repeatability of the adaptive path taken from the same starting point (see Box 1 in Venkataram and Kryazhimskiy (2023) for definitions). The contrasting effects of intragenomic epistasis on these two forms of repeatability are not necessarily in contradiction. Decreased endpoint repeatability could result from increased path repeatability when highly conserved paths from different initial starting points lead to distinct peaks of a multi-peaked landscape. In experimental evolution, path repeatability has often been addressed by evolving replicates of populations from the same starting genotype (e.g., Good et al., 2017). Conversely, endpoint repeatability could be captured by evolving populations starting from multiple different genetic backgrounds (e.g., in an evolution replay experiment; Decaestecker et al., 2007).

In contrast to intragenomic epistasis, intergenomic epistasis reduces the number of adaptive outcomes, even in rugged landscapes, as not all peaks are joint fitness peaks. So, although intergenomic epistasis generates additional peaks in the metagenotype space similar to intragenomic epistasis (resulting in more possible outcomes of adaptation across coevolutionary fitness landscapes), the species interaction simultaneously limits the number of possible adaptive equilibria (fewer realized outcomes of adaptation per coevolutionary fitness landscape). This led to high endpoint repeatability at the level of individual landscapes, but increased variability between different landscapes generated with the same parameters. Additionally, we the absence of a joint fitness peak can result in cycling between metagenotypes. Capturing the observed coevolutionary constraint on adaptation imposed by intergenomic epistasis in an empirical system would presumably require detailed knowledge of the fitness mappings of the interacting species. One approach to infer such information is via so-called coevolutionary genome-wide association studies (co-GWAS), which have been developed to identify genetic polymorphisms between species and link them to fitness or phenotypes (Ebert & Fields, 2020; MacPherson et al., 2018; Märkle et al., 2021; Nuismer et al., 2022).

The question of evolutionary repeatability and the study of epistasis often go hand in hand with the concept of evolutionary predictability (de Visser & Krug, 2014; Johnson et al., 2023; Wortel et al., 2023). Repeatability usually increases predictability, but here we find that our results, though repeatable, could be difficult to predict. In our model, the location of joint fitness peaks on the coevolutionary fitness landscapes is an emergent feature that could not be captured by global properties of individual fitness landscapes, such as the number of genetic interactions per locus (*K* and *C*) or the landscape ruggedness. Furthermore, landscape metrics such as the inferred correlation of fitness ranks between species and ruggedness could not explain the fitness ranks of observed adaptive outcomes for individual species or the occurrence of cycling, even though landscape ruggedness is expected to be a determinant of the probability of reaching high fitness peaks (Li & Zhang, 2025). It is likely that other global summary statistics based on ruggedness are similarly insufficient to capture the complexities of differing genetic interactions and fitness mappings (Nowak & Krug, 2015).

Does coevolution help or hinder a species’ or a community’s adaptation? To answer this question, we quantified the individual and joint fitness load of two interacting species at the end of adaptation. Instead of adaptation being skewed towards higher fitness fitness peaks as in the *NK* model, or (even more so) on empirical fitness landscapes like the *Escherichia coli* dihydrofolate reductase gene *folA* (Hunter & Martin, 2026; Li & Zhang, 2025; Papkou et al., 2023), we saw many coevolutionary fitness landscapes where adaptive walks repeatedly led to low fitness endpoints. With intergenomic epistasis, we observed a pervasive coevolutionary fitness load that was only partially explained by whether the species interaction, inferred from the correlation of fitness ranks between species, was antagonistic or synergistic. A larger fitness load was associated with stronger negative fitness rank correlations and thus more antagonistic interactions between species. This aligns with our ecological understanding of systems with competitive, host-parasite, or predator-prey relationships, where adaptation of one species is constrained by its interaction with another. For example, developing and maintaining immunity to parasites can be costly as it diverts resources from reproduction (De Lisle & Bolnick, 2021; Lochmiller & Deerenberg, 2000). Interestingly, the fitness load caused by intergenomic epistasis was consistently larger than that in a corresponding “seascape” model in which a single species adapts to a fluctuating environment.

Our results show that coevolutionary landscapes generate coevolutionary load even when the fitness ranks are positively correlated. Notably, this does not have to be at odds with our understanding of mutualisms, as they often involve a delicate balance between the costs and benefits of an interaction (Bronstein, 2001; Heath, 2010). An intuitive example of this is found in the plant-pollinator world. Many flowering plants are dependent on pollinators for successful reproduction, but might produce costly nectar in order to attract beneficial pollinators (e.g., Southwick, 1984). In such systems, successful adaptation may be less about whether evolution has had time to take species to a place on the fitness landscape that maximises fitness for both interacting species, and more about whether such a joint global fitness peak even exists given inherent physiological and genetic limitations.

An outcome unattainable in most classical fitness landscape models is cycling. As already described in Kauffman and Johnsen (1991) and Bak et al. (1992), we recovered that cycling between metagenotypes became more common with increasingly prevalent intergenomic interactions. Notably, in the *NKC* model, cycling occurred in the absence of any population dynamics and resembles an ongoing arms race through reciprocal beneficial substitutions between hosts and parasites (Wilfert & Jiggins, 2013). We found that cycling outcomes overall showed increased fitness loads for both species and communities, making cycling a (from a fitness-maximization viewpoint) undesirable outcome. However, the fitness load difference between species was lower than average when cycling outcomes were observed. Thus, cycling might strengthen or stabilize coexistence between species in the long term, even when interactions are antagonistic.

In our simulations, we found that the way fitness landscapes were navigated (e.g., the type of adaptive walk or asymmetry in the mutation rate between the species) did not affect endpoint repeatability. These almost-deterministic results might be a by-product of studying small landscapes. This choice of parameter space might affect the observed role of different adaptive walks in creating path dependencies and branching points along the way. For larger landscapes, we might recover the results of Bull (2021) and Hordijk and Kauffman (2005), who found that taking multiple adaptive steps at a time was beneficial for species adapting on larger *NKC*(*S*) landscapes with *N* = 100 loci.

In general, restricting this study to small fitness landscapes allowed us to analyze the differing effects of intra- and intergenomic epistasis in detail and study how they affect the number of realized adaptive outcomes. It helped limit the number of parameters to study and allowed for the dissection of the effect of all possible interaction structures. However, future work should address how our results hold in larger, more complex coevolutionary fitness spaces. For example, it will be interesting to study whether cycling becomes more or less common on larger fitness landscapes.

Throughout the text, we compared coevolutionary *NKC*(*S*) landscapes with *N* = 2 loci to single-species *NK* landscapes and *NK*(*E*) seascapes with *N^′^* = 4 loci, since they represent a similar parameter space in terms of number of genetic interactions, landscape ruggedness, and number of (meta)genotypes. However, they differ in their connectivity between (meta)genotypes, which is crucial for determining how the species can navigate their fitness landscape. In practice, this means that though the *NKC*(*S*) and the *NK*(*E*) model both share their multidimensional fitness mapping, in the *NKC*(*S*) model at each step only *N* loci can mutate, whereas in the *NK*(*E*) model, all *N^′^* = *N* + *N* loci can mutate. Thus, species on *NK*(*E*) seascapes have access to a larger pool of potentially beneficial mutations and therefore a larger evolutionary potential. This difference might also explain why we found an effect of ecological interactions on endpoint repeatability, whereas Amado and Bank (2024), who studied interactions between genotypes of the same species during polymorphic adaptation, did not.

Our model assumes strictly monomorphic populations of essentially infinite size moving through genotype space in a step-wise fashion. Introducing explicit population dynamics and genetics on coevolutionary fitness landscapes could allow for population diversification and extinction, which are not considered as possible events in this study. This model complexification would also allow us to study other aspects of a species’ evolutionary potential, such as the importance of absolute mutational input, differing population sizes (e.g., differing levels of genetic drift), and species loss (Gandon & Michalakis, 2002; Szendro, Franke, et al., 2013). One way to implement density-dependent dynamics is to link the *NKC*(*S*) model to Lotka-Volterra population dynamics, as done in the original paper by Kauffman and Johnsen (1991). Here, fitness and interaction coefficients were assigned to each genotype to model population abundances. This would make the model more ecologically explicit and relieve the assumptions of strong selection and weak mutation.

The *NKC*(*S*) model is a probabilistic fitness landscape model, which, on the one hand, helps build landscapes with few parameters, and on the other hand, makes it difficult to interpret the generated landscapes with respect to any mechanistic features of real interacting organisms. To generate more easily interpretable coevolutionary fitness landscapes, one could implement specific genetic interactions with explicit epistasis terms to investigate how repeatability varies across different genetic interaction networks. One way to do this is with the model proposed in Hablützel and Bank (2025), which allows one to individually specify additive and epistatic terms of intra- and intergenomic interactions, including pairwise and higher-order epistatic terms, to emulate well-known genetic interaction patterns such as gene-for-gene interactions or allele-matching. It would be interesting to use such a model to test whether it recovers the interaction network-specific patterns of adaptation found in the present study.

In summary, we found that repeatability can depend strongly on the genetic architecture of interacting species, even in the absence of stochastic mutation or population dynamics. Furthermore, intergenomic epistasis caused highly repeatable patterns of adaptation that deviated from expectations based solely on intragenomic epistasis due to the frequent emergence of fitness trade-offs between species. Understanding when and where such fitness trade-offs emerge requires knowledge of the given coevolutionary fitness landscape; assessing the fitness landscapes of one of the interacting species is insufficient. This underlines the importance of explicitly considering interactions between species and their consequences on adaptation in the study of ecological and evolutionary dynamics.

## Data availability

All annotated code to reproduce this study can be found under https://github.com/banklab/Loraine-NKCS and will be deposited on Zenodo upon acceptance of the paper. Supporting model information can be found in Supplement A and supplementary figures in Supplement B.

## Acknowledgements

We thank the THEE division and the ComBi group for helpful discussions. We thank three anonymous reviewers for their careful reading of the manuscript and their helpful comments and suggestions. Simulations were performed on UBELIX (https://www.id.unibe.ch/hpc), the HPC cluster at the University of Bern.

## Supplement A: Additional model specifications

### Construction of the NKC(S) landscapes

We construct the *NKC*(*S*) fitness landscapes using three building blocks: (i) a table of all possible metagenotypes in the system, (ii) interaction matrices linking the epistatically interacting loci to each other (a matrix for intragenomic epistasis and a matrix for intergenomic epistasis for each species), and iii) a table with all possible fitness contributions depending on the different genetic backgrounds. Here we describe how to obtain these building blocks for a two-species community.

### Metagenotypes

A system with two haploid species with *N* loci and *A* allelic variants has *A^N^*^+^*^N^* unique metagenotypes{⃗F, P⃗ } containing all possible allelic combinations. A metagenotype is the combination of genotypes of the focal and partner species in the community.

### Interaction matrices

The interaction matrices (IMs) describe the epistatic interactions among loci. Within a species, the fitness contribution 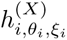 at locus *i* depends on *K* + 1 loci - the allele present at locus *i* itself and the alleles present at *K* other loci. In the IM, an interaction between two loci is represented by a 1, and the absence of an interaction between two loci is represented by a 0. For *N* = 2, intragenomic epistasis is always reciprocal, and the possible interaction matrices either take the shape

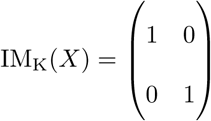

in the absence of intragenomic epistasis (i.e., the fitness contribution 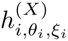 at locus *i* only depends on the allele at locus *i*; *K* = 0), or

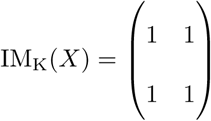

in the presence of intragenomic epistasis (i.e., the fitness contribution 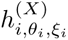 at locus *i* depends on the alleles present at both loci in the genome; *K* = 1).

However, for *N* = 2 and *C* = 1, genetic interactions between genomes are not necessarily reciprocal. Here, IM_C_(F, P) describes which loci in the genome of species P affect which loci in the focal species F. This could, for example, yield

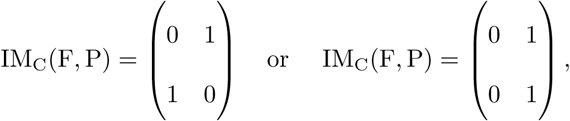

resulting in different possible interaction networks between the genomes. Here, we investigate IMs that differ in their symmetry between the species and their specificity of interactions between loci. Therefore, we implement the four possible interaction types based on the combination of different interaction matrices for *NKC*(*S*) landscapes with *C* = 1. Especially for larger genomes (i.e., more loci), the type of interaction matrix could play a role in mediating genetic interactions. I.e., there we can consider matrices where interacting loci are distributed randomly in the genome, or arranged in specific patterns, e.g. in blocks of interacting loci, or following a nearest-neighbour or preferential attachment method (see Nowak and Krug, 2015; Rivkin and Siggelkow, 2007; Welch and Waxman, 2005). We explore a selection of such scenarios for intragenomic interactions for *NK* landscapes with *N^′^* = 4 loci in Supplement B .

**Table A.1:**
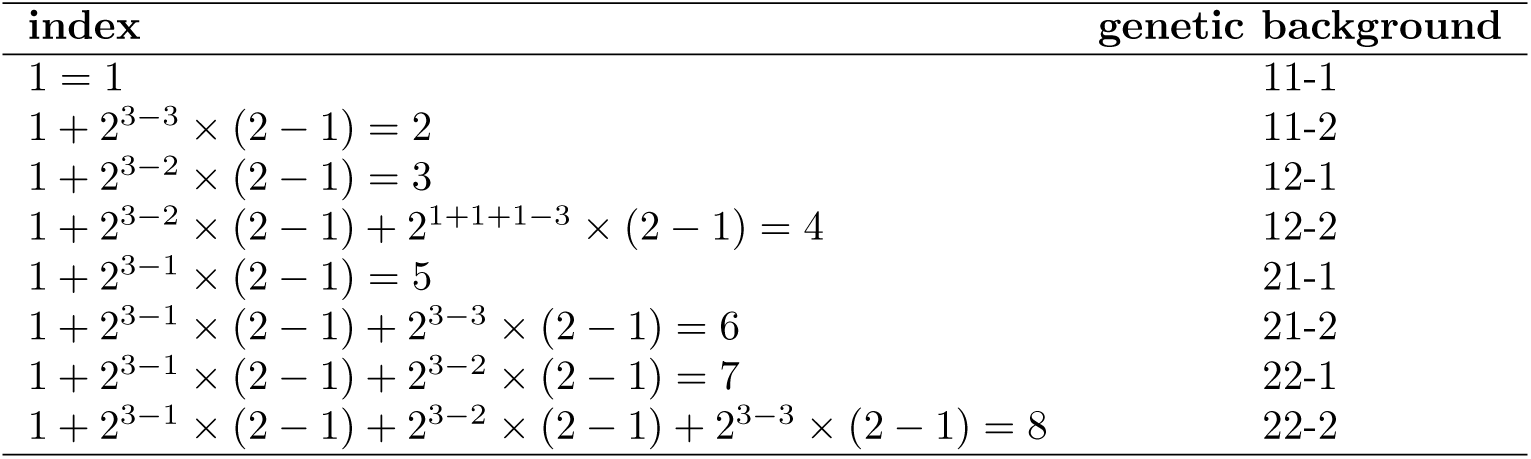
Example for how to index the rows in a lookup table for *N* = 2, *A* = 2, *K* = 1, *C* = 1, *S* = 1. We calculate the index following eq. (A.1). The total number of rows is *A^K^*^+^*^C^*^+1^ = 8 and each index is unique for a given genetic background, where we are only looking at the allelic states of the *K* + *C* + 1 loci of interest. The genetic background is given by the alleleic states of 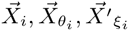 translated into *σ_j_ ∈ {*1, 2*}* for reference and mutant allele.

### Look-up table for fitness contributions

For each species, there are *A^K^*^+(^*^S×C^*^)+1^ possible fitness contributions 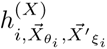 at a locus *i* depending on the relevant allele combinations for all loci 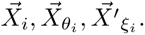 For each species *X*, we construct a table with *N* columns (= all the loci of a species) where each row represents a given genetic background (*A^K^*^+(^*^S×C^*^)+1^ possible backgrounds). This way, we can access the fitness contribution 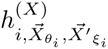 at each locus *i* given its genetic background. Accessing the correct value works by translating the genetic background we are interested in into a unique index of the look-up table. For this, we iterate over all *K* + (*S* × *C*) + 1 loci of interest that affect the fitness contribution at a given locus *i* and read out their allelic values. We calculate the row index as:

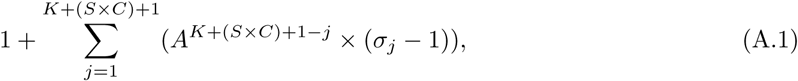

where *j* ∈ {1, 2*, ..K* + (*S* × *C*) + 1} is a locus of interest in 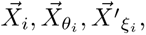 and *σ_j_* is the allele present at the locus of interest. For this we translate alleles into numbers *σ_j_* ∈ {1*, …, A*}, where the reference (wildtype) allele has value 1 and the other alleles take on values 2*, …, A*. See Table A.1 for how to calculate indices to access fitness contributions 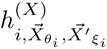 in the above described look-up table.

### Generating the fitness landscape

Using the interaction matrices, we identify the loci of interest (= the *K* + *C* interacting loci and the focal locus itself) for each focal locus *i*. For each metagenotype {⃗F, P⃗ }, we then extract the allelic values at the loci of interest, based on which we find 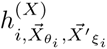 in the look-up table. This yields the fitness contributions 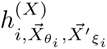 at each locus *i* ∈ {1*, …N*} in the focal genome. Now, we calculate fitness *W* (*X* | {⃗F, P⃗ }) for each metagenotype {⃗F, P⃗ } and each species *X* (*X* ∈ {F, P}) using eq. (1).

Thus, putting everything together, we generate a full table of metagenotypes, the resulting fitness contributions 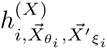 at each of the *N* focal loci and the resulting fitness *W* (*X* | {⃗F, ^P⃗^}). This results in a fitness landscape for each species, where fitness can vary for the same genotype *X⃗* of a species due to intergenomic epistasis if *C* ≠ 0, when the metagenotype {⃗F, P⃗ } differs.

### Construction of the *NK*(*E*) model

Our implementation of the *NK*(*E*) model model is analogous to the *NKC*(*S*) model explained in the main text in many aspects. However, here we model a single species that experiences *E* different environments, where in each environment *Y*, a different trait is under selection. The species therefore has *E* environment-associated traits, each encoded by *N* loci, that confer the species environment-specific fitness *W* (F | F⃗*, Y*). (Note that we are assuming absence of pleiotropy throughout, but an extension to varying degrees of pleiotropy is straightforward to imagine.) Practically, there are three main differences between the *NK*(*E*) implementation and a regular *NK* model: (i) the structure of the intragenomic interaction matrix, (ii) how we calculate environment-specific fitness, and (iii) that we switch between which trait (set of loci) is under selection (i.e., contributes to fitness) as we are switching between environments. Table A.2 explains the main changes in notation and figure A.1 illustrates the *NK*(*E*) model. Additionally, we highlight the main points of comparison between the *NK*(*C*) and the *NK*(*E*) model in Fig. A.2.

### Interaction matrices

For the *NK*(*E*) model, all genetic interactions happen within the same genome (intragenomic epistasis), however we structure the genome into *E* blocks of *N* loci each to encode different environment-associated traits. Therefore, we can distinguish between genetic interactions of loci within the same trait, or between different traits, which we denote by *K_w_* and *K_b_*, respectively. The resulting interaction matrices are modular in nature. For example, for *N* = 2, *E* = 2, *K_w_* = 1 and *K_b_* = 0, the resulting interaction matrix takes the following form:

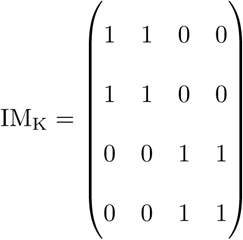

As for the *NKC*(*S*) model, *K_b_* can be arranged differently regarding symmetry between traits. For this we incorporate the same interaction matrices as in the *NKC*(*S*) model (see Fig. 6) into the off-diagonal blocks.

**Table A.2:** Equivalence of notation between the *NKC*(*S*) model and the *NK*(*E*) model. Instead of different species, we model different environment-associated traits encoded by *E* blocks of *N* loci in a single species’ genome.

| Description | $NKC(S)$ | $NK(E)$ |
| --- | --- | --- |
| The context on which fitness depends on | species $X$ | environment $Y$ |
| The unit of selection | species-level | trait-level |
| The number of contexts to consider per species | $S$ partners | $E$ environments |
| The number of loci in a species' genome | $N$ | $N' = E \times N$ |
| Genetic interactions between loci within the same unit of selection (species/trait) | $K$ | $K_w$ |
| Genetic interactions between loci between different units of selection (species/trait) | $C$ | $K_b$ |
| Fitness contribution of locus $i$ in a species | $h_{i, \vec{X}_{\theta_i}, \vec{X}'_{\xi_i}}^{(X)}$ | $h_{i, \vec{F}_{\theta_i}, \vec{F}'_{\xi_i}}^{(Y)}$ |
| The set of indices of loci in the same unit of selection (species/trait) interacting with locus $i$ | $\theta_i$ | $\theta_i$ |
| The set of indices of loci a different unit of selection (species/trait) interacting with locus $i$ | $\xi_i$ | $\xi_i$ |
| Number of mutations possible at a time per context (species/environment) | $R$ | $R$ |

**Figure A.1:**
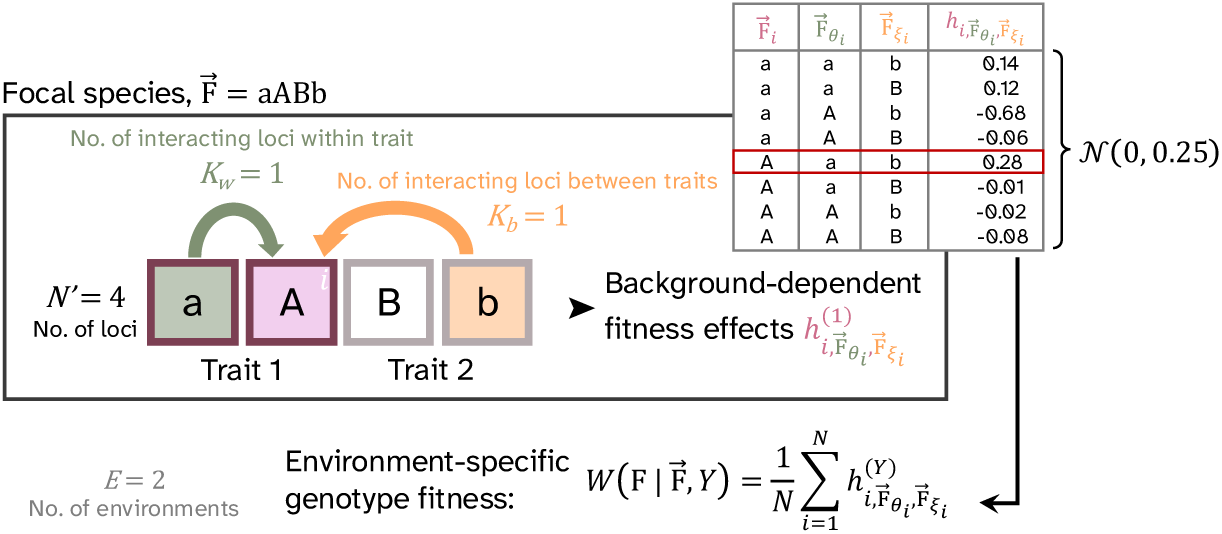
A visual representation of how to compute the fitness contribution of a locus *i* in the *NK*(*E*) model. For each environment *Y*, there is a trait encoded by *N* = 2 loci with *A* = 2 alleles each. Alleles are encoded as F⃗*_i_* ∈ {a, A} for the loci in trait 1 and F⃗*_j_* ∈ {b, B} for the loci in trait 2. Each locus *i* (here *i* = 2 is highlighted in pink) interacts with *K_w_* loci in its own trait (green) and *K_b_* loci in other traits (orange). Depending on the combination of alleles at all relevant loci 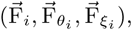 the fitness contribution of locus 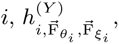 differs. There are 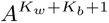 possible values for 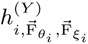 where each value is drawn from a normal distribution and associated with a different genetic background (see top right table). The fitness of a genotype species is then calculated in a given environment as the average of all *N* fitness contributions in the relevant environment-associated trait.

### Calculating fitness

In the *NK*(*E*) model, fitness is mediated by environment-specific traits; e.g. imagine a population of soil bacteria living in a river bank that frequently gets flooded changing the oxygen levels and flushing out nutrients in the soil. I.e., in a given environment, fitness only depends on the respective environment-associated trait. Fitness of our focal species *W* (F | F⃗*, Y*) thus depends on its genotype F⃗ and the environment *Y* . Fitness is calculated based on the fitness contributions 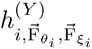 of the *N* loci in the environment-associated trait, their individual allelic states F⃗*_i_* and the allelic states at the loci that interact within the same trait 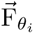 and the loci that interact between different traits 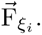

**Figure A.2:**
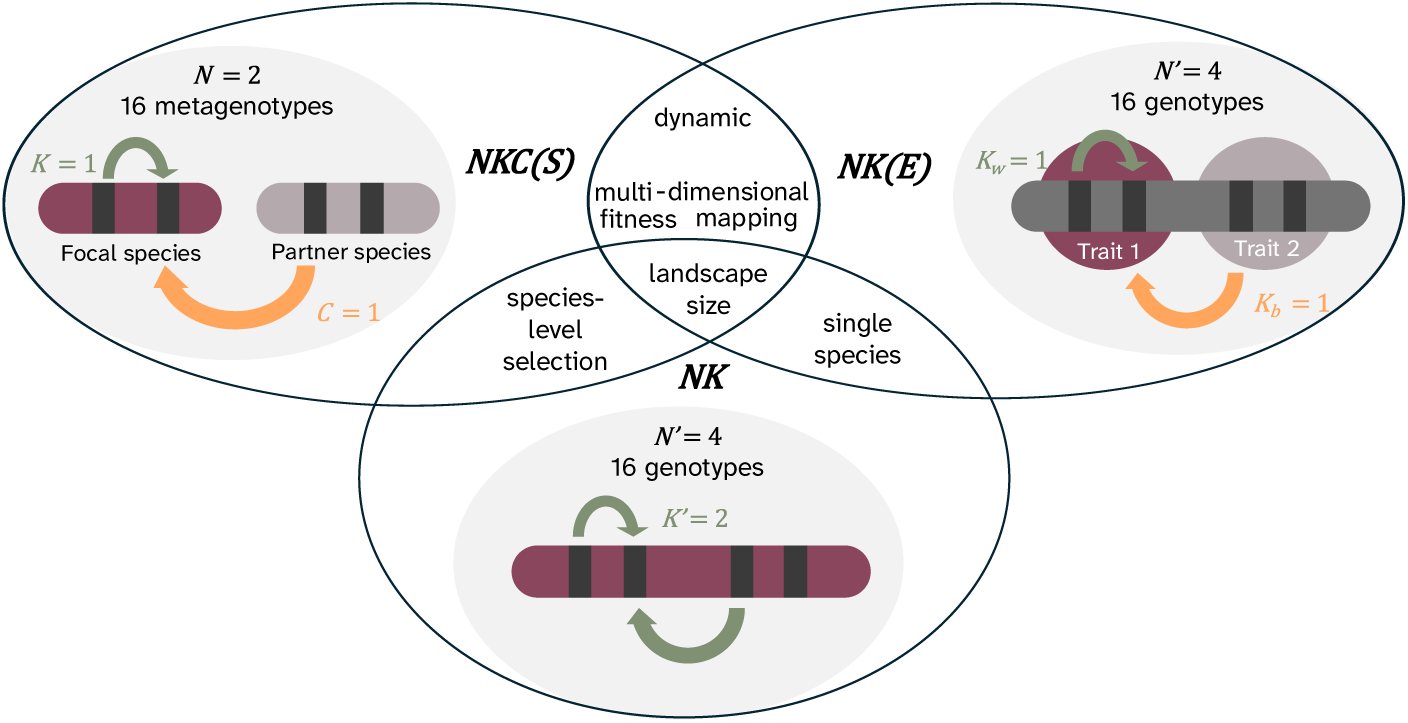
Differences and commonalities between the three fitness landscape models we investigate here. The *NKC*(*S*) and the *NK*(*E*) model both generate dynamic fitness landscapes, where fitness of a (meta)genotype can either differ between species (same metagenotype in different species = different fitness in the *NKC*(*S*) model), or between abiotic environments (same genotype in different environment = different fitness in the seascapes model).

### Switching environments

Depending on the environment, we switch between which loci are evaluated for fitness. This switch between environments is determined by the rounds of (possible) mutations *R* the species spends in each environment and can be implemented in three ways; (i) switch after each (possible) mutation step (i.e, regular switching between environments (*R* = 1)), (ii) spending more time in one environment than the other (i.e., regular switching but with asymmetry between environments *R_Y_* ; *R*_1_ = 2*, R*_2_ = 1), (iii) irregular switching between environments, where the change rate between environments was drawn from a Poisson distribution (*R_Y_* ∼ Pois(1)).

### Adaptive walk algorithms

When simulating adaptive walks we implement a strong selection weak mutation (SSWM) regime following three possible fitness-increasing algorithms: (i) random walks randomly chose any (meta)genotype of higher fitness that is one mutational step away from the current (meta)genotype. In the *NKC*(*S*) model this only includes metagenotypes that can be reached through mutations in the genome of whichever species is currently mutating. (ii) True adaptive walks pick higher-fitness single-step mutational neighbours proportional to their fitness increase. I.e., adaptation is skewed toward larger fitness-increasing steps. (iii) Greedy walks always pick a mutational neighbour that maximises the fitness increase of the adaptive step. Throughout the main text all results were generated using random adaptive walks, unless explicitly stated otherwise.

## Supplement B: Supplementary figures and results

### Comparing landscape ruggedness

To estimate the ruggedness of our fitness landscapes, we compared how strongly the fitness mappings of our generated fitness landscapes deviate from a linear approximation. For this, we fitted linear models to the landscapes to calculate the slopes of each fitness landscape as the mean change in fitness by change in allelic value, and roughness as the variance of residuals of each linear model. We then use the ratios of roughness to slope as estimates of landscape ruggedness (Bank et al., 2016; Szendro et al., 2013). For fully additive landscapes, we expect the fitness mapping to be linear and the ruggedness to be 0. With epistasis in the landscape, the roughness to slope ratio increases *>* 0 due to deviations from the linear model and thus indicate ruggedness of the landscape.

**Figure B.1:**
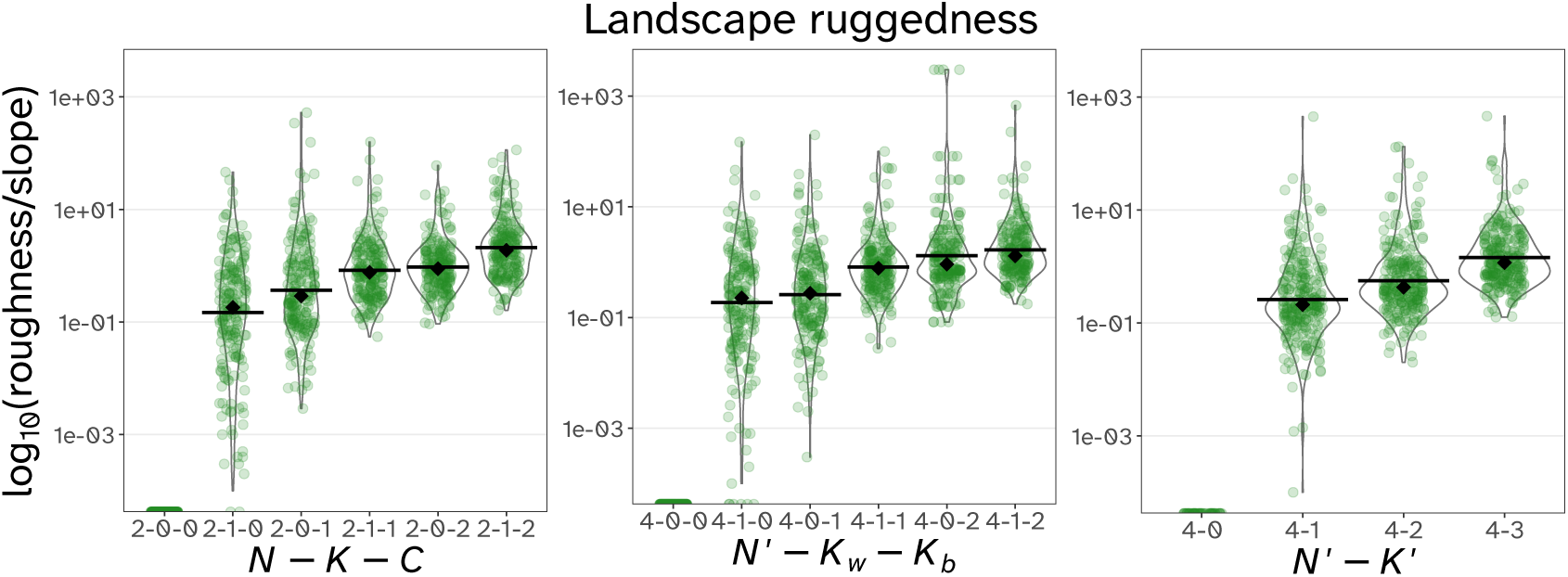
Measured roughness to slope ratio as an estimate for landscape ruggedness. For all three models treated in the main text and supplementary materials, we show the ruggedness of 200 land- or seascape replicates, with their overall means (black bars) and medians (black diamonds). Ruggedness increases with the number of genetic interactions per locus (*K*, *C*, *K_w_* or *K_b_*). Land- and seascapes with *N^′^* = 4 show comparable levels of ruggedness to landscapes with *N* = 2 and different levels of epistasis.

### The effect of interaction matrices on adaptive outcomes

**Figure B.2:**
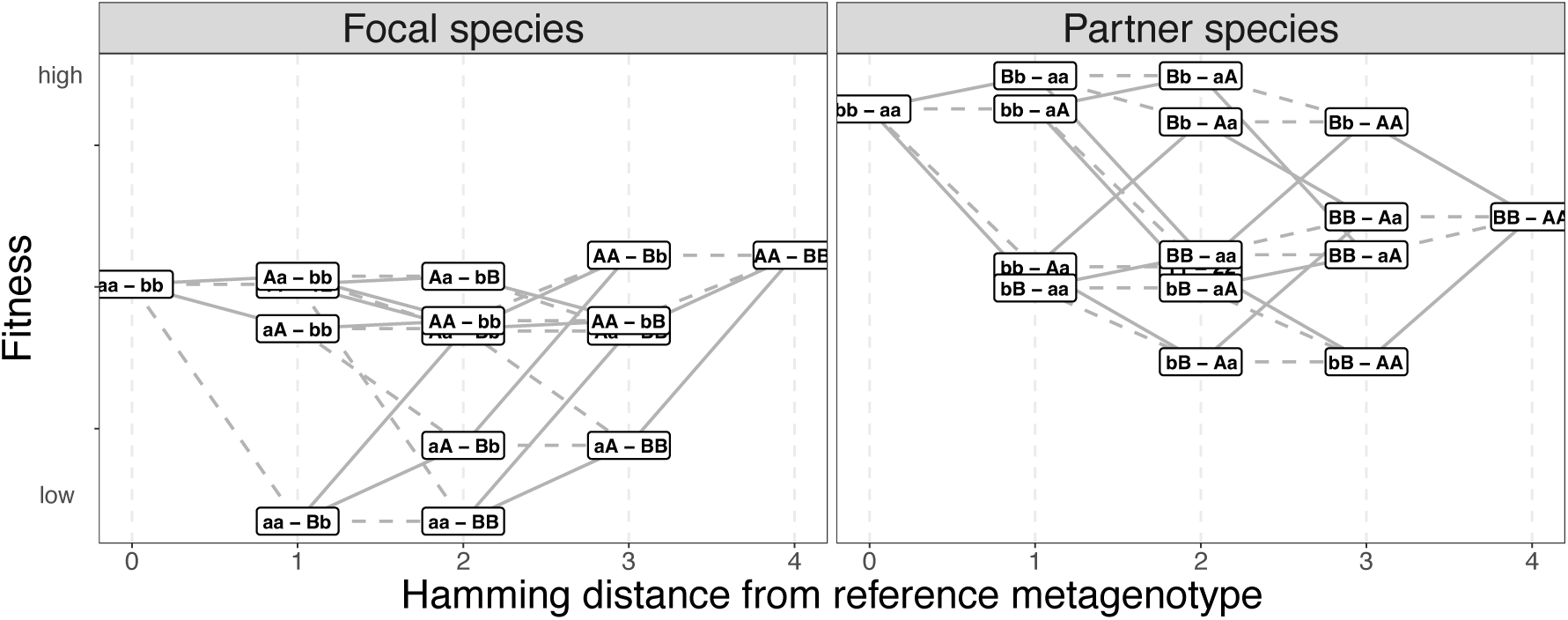
Interaction network 1 generates neutral plateaus in the fitness mapping by introducing redundancy between some of the metagenotypes; the way this manifests on a species’ fitness landscape is by introducing parallel congruent sublandscapes. Example of a pair of fitness landscapes generated with *N* = 2*, K* = 0*, C* = 1*, A* = 2. Alleles are encoded as F⃗*_i_* ∈ {a, A*}* and P⃗*_i_* ∈ {b, B} and the reference point (Hamming distance 0) is {F⃗, P⃗ } = (aa, bb). Solid (dashed) lines show mutational steps in the focal (partner) genome.

**Figure B.3:**
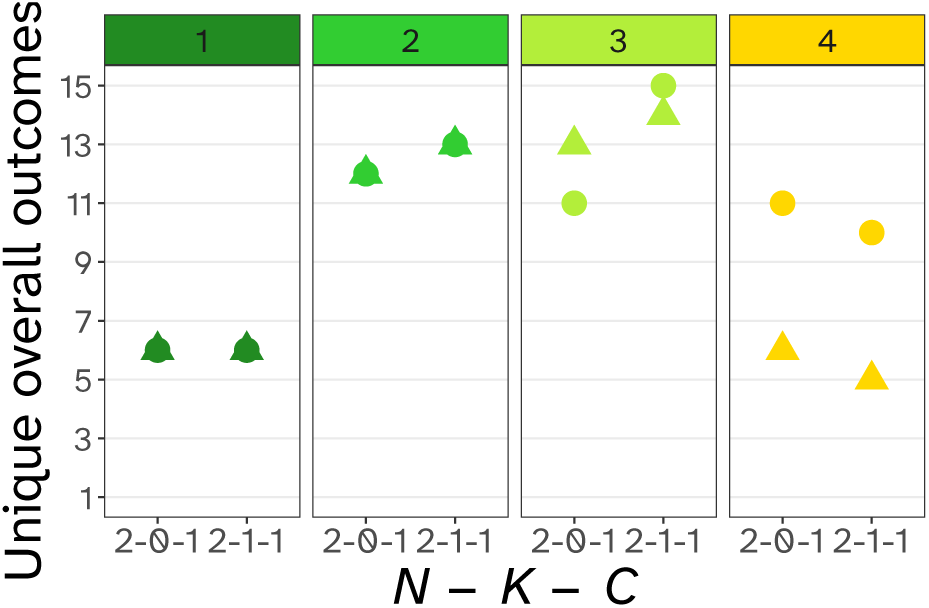
The total number of unique outcomes observed differs across the four investigated interaction matrices. For each parameter combination with *C* = 1 we count the unique outcomes for both interacting species (circle and triangle). A low number of unique outcomes leads to low variability across different fitness landscape replicates. See Fig. 6 for the explanations of interaction matrices.

**Figure B.4:**
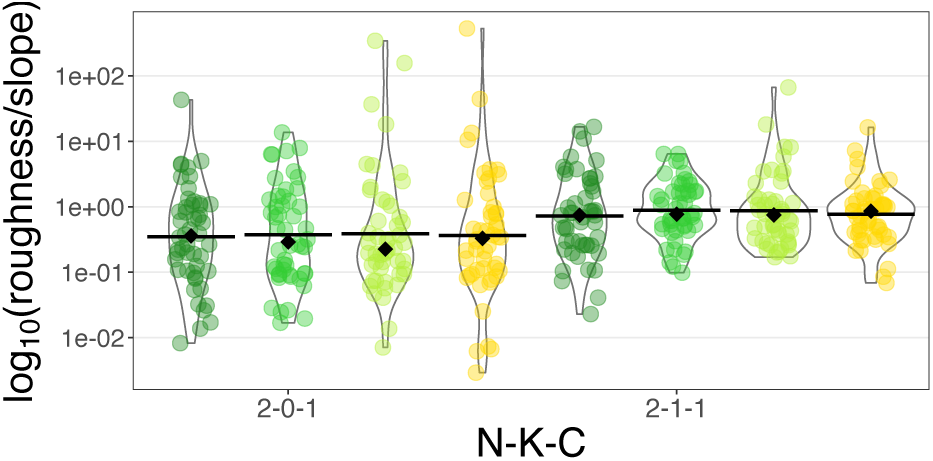
Measured roughness to slope ratio as an estimate for landscape ruggedness. Here we compare the ruggedness of different landscapes generated with different interaction networks for *C* = 1. There is no major difference in the ruggedness. We show mean (black bar) and median (black diamond) ruggedness. For explanations of the different interaction networks, see Fig. 6.

**Figure B.5:**
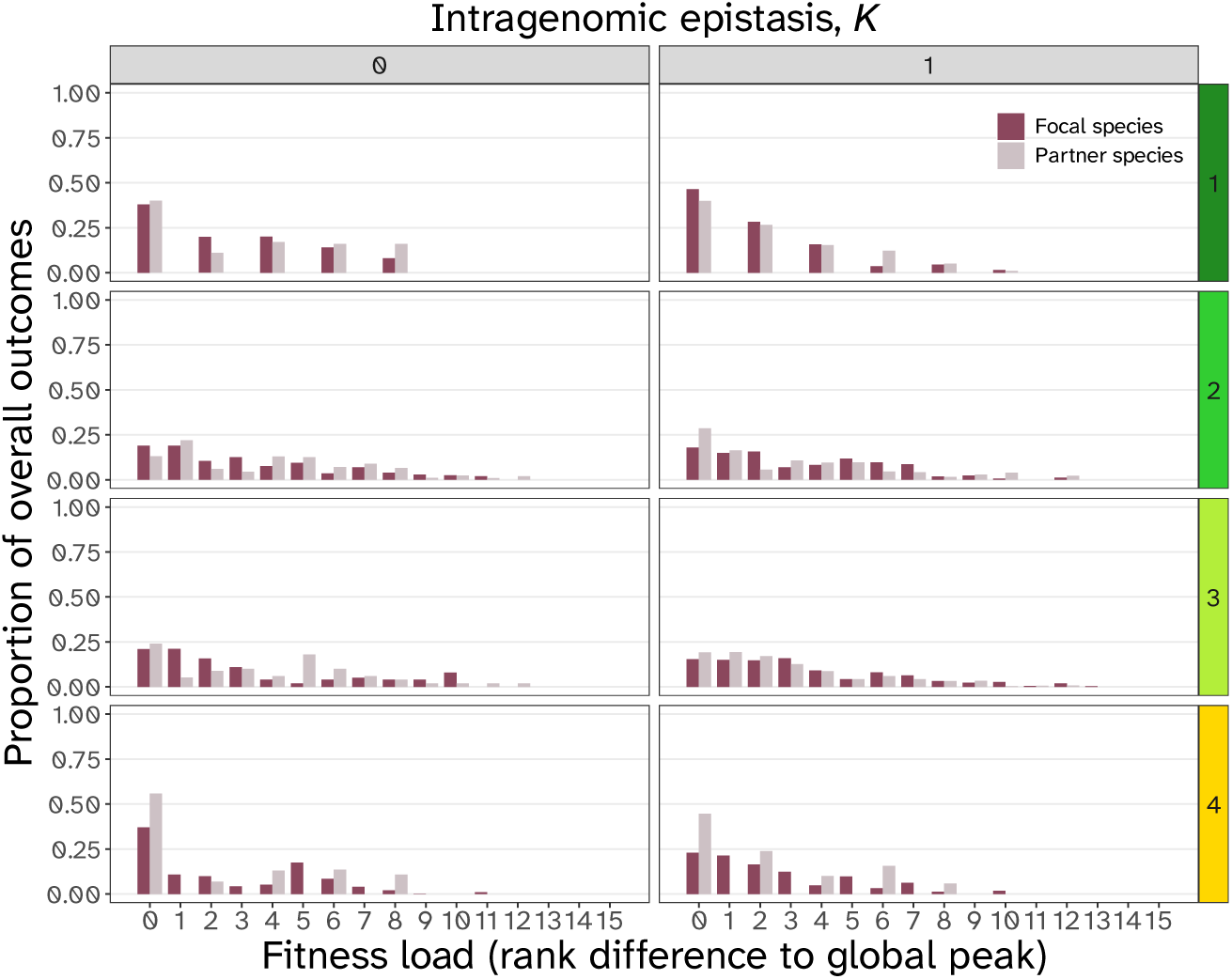
The coevolutionary fitness load depends on the underlying interaction network for *C* = 1 and can differ between species. The fitness load was reduced for species with specific genetic interactions (interaction networks 1 and 4). Here, we included cycling outcomes by calculating the mean fitness of a species during the cycle. Explanations of the interaction networks can be found in Fig. 6.

### Adaptive walk types have little effect on repeatability and variability

Here, we report results from greedy or true adaptive walks on the same landscapes described in the main text. Notably, cycles persist under different kinds of adaptive walks and differences between the adaptive walk types are marginal.

**Figure B.6:**
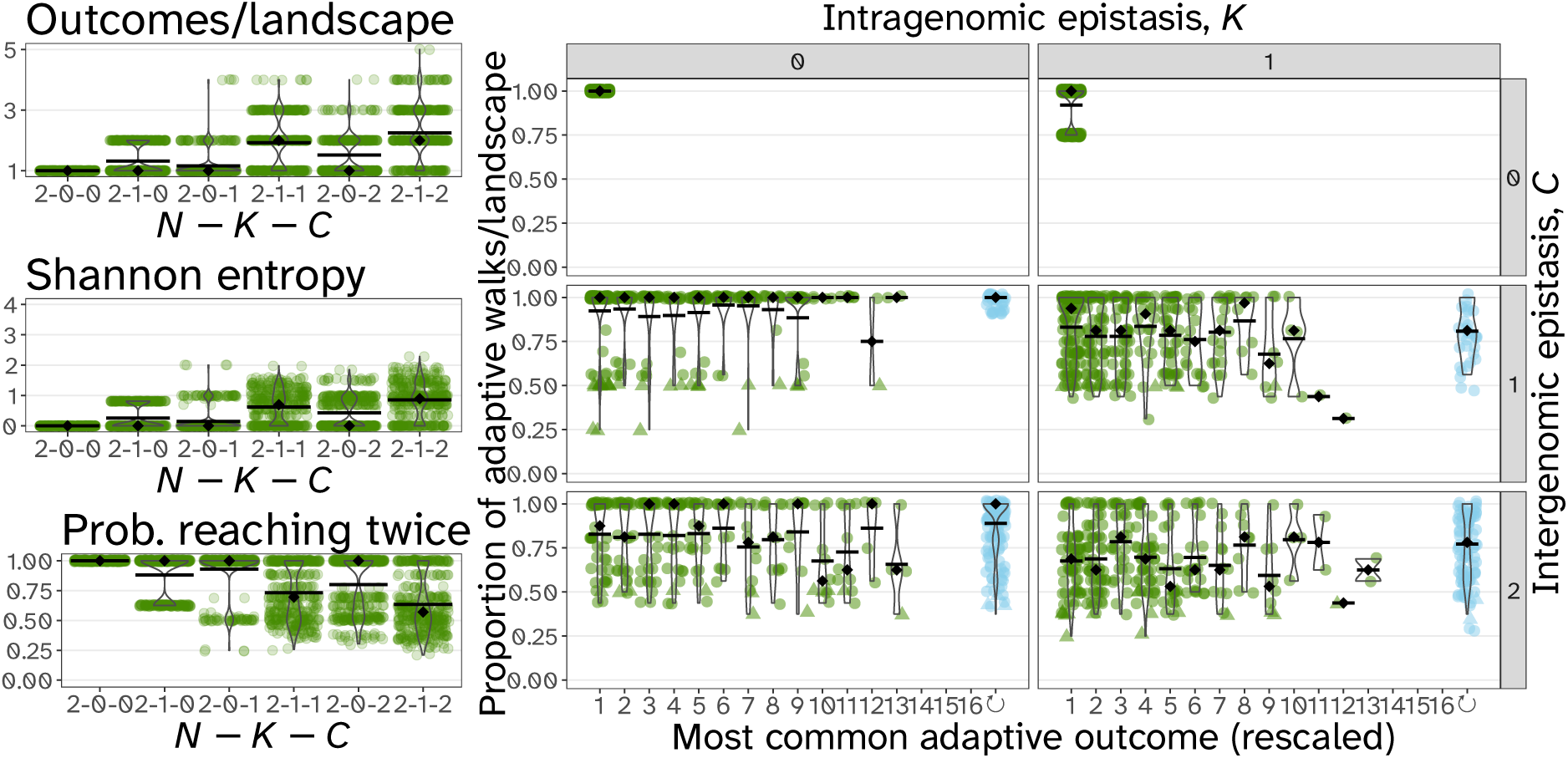
Repeatability results for greedy walks for *N* = 2 and variable combinations of *K* and *C*. We observe the same main patterns as in Fig. 4a,d,g and 5a.

**Figure B.7:**
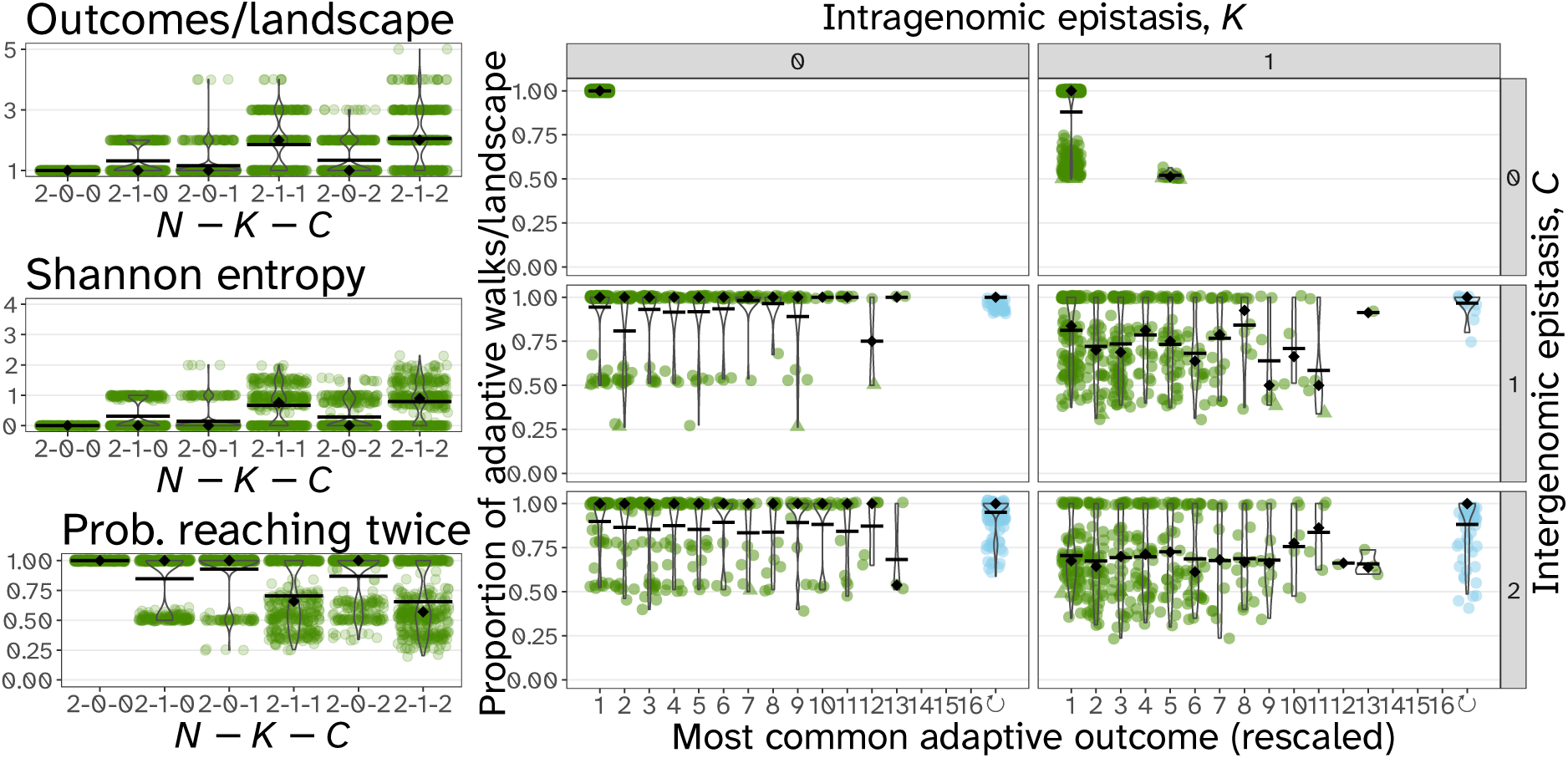
Repeatability results for true adaptive walks for *N* = 2 and variable combinations of *K* and *C*. We observe the same main patterns as in Fig. 5a.

**Figure B.8:**
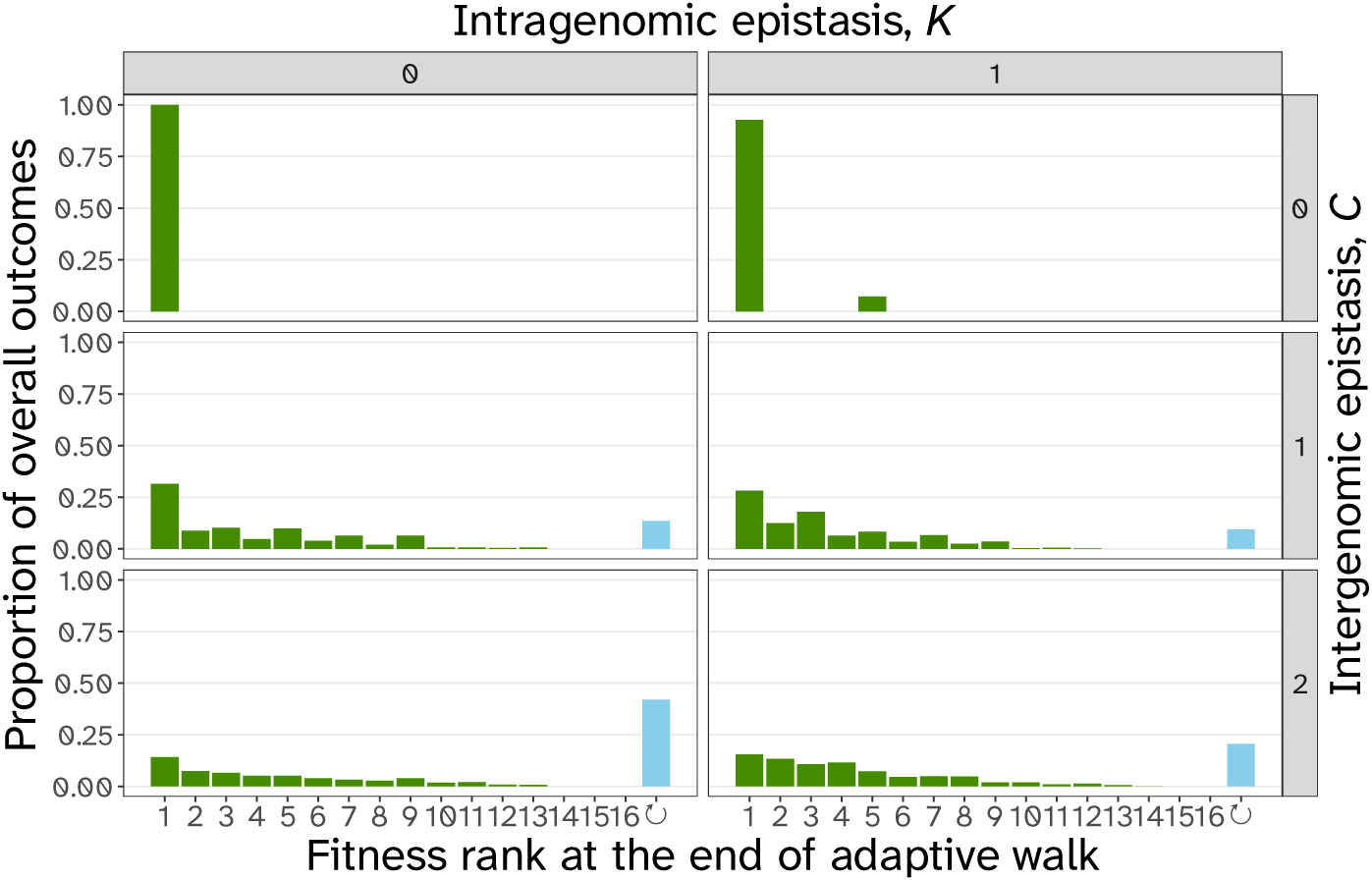
Distribution of adaptive outcomes for *N* = 2 for greedy adaptive walks. We observe the main patterns as in Fig. B.10.

**Figure B.9:**
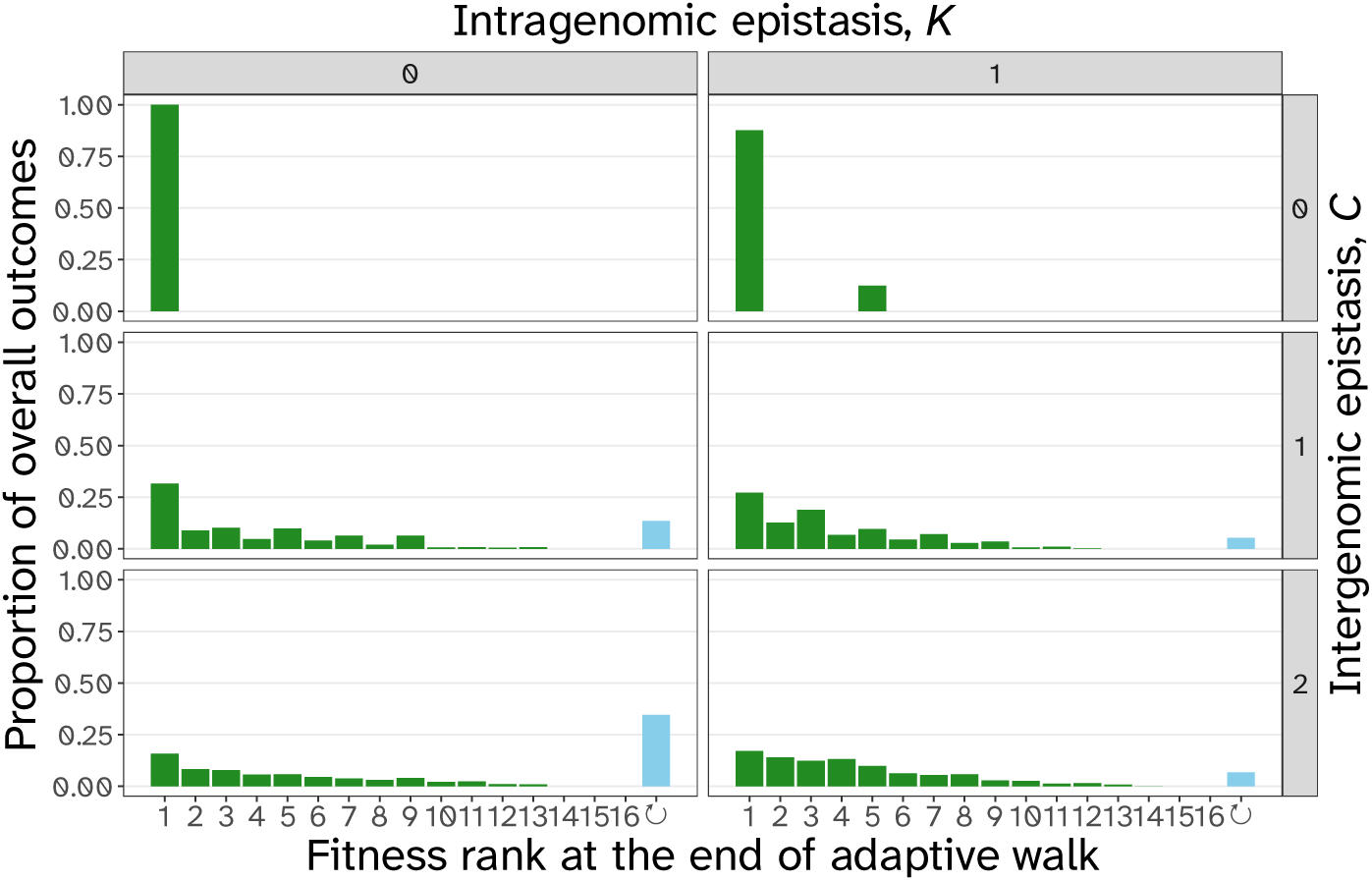
Distribution of adaptive outcomes for *N* = 2 for true adaptive walks. We observe the main patterns as in Fig. B.10.

### Fitness load for species and communities

**Figure B.10:**
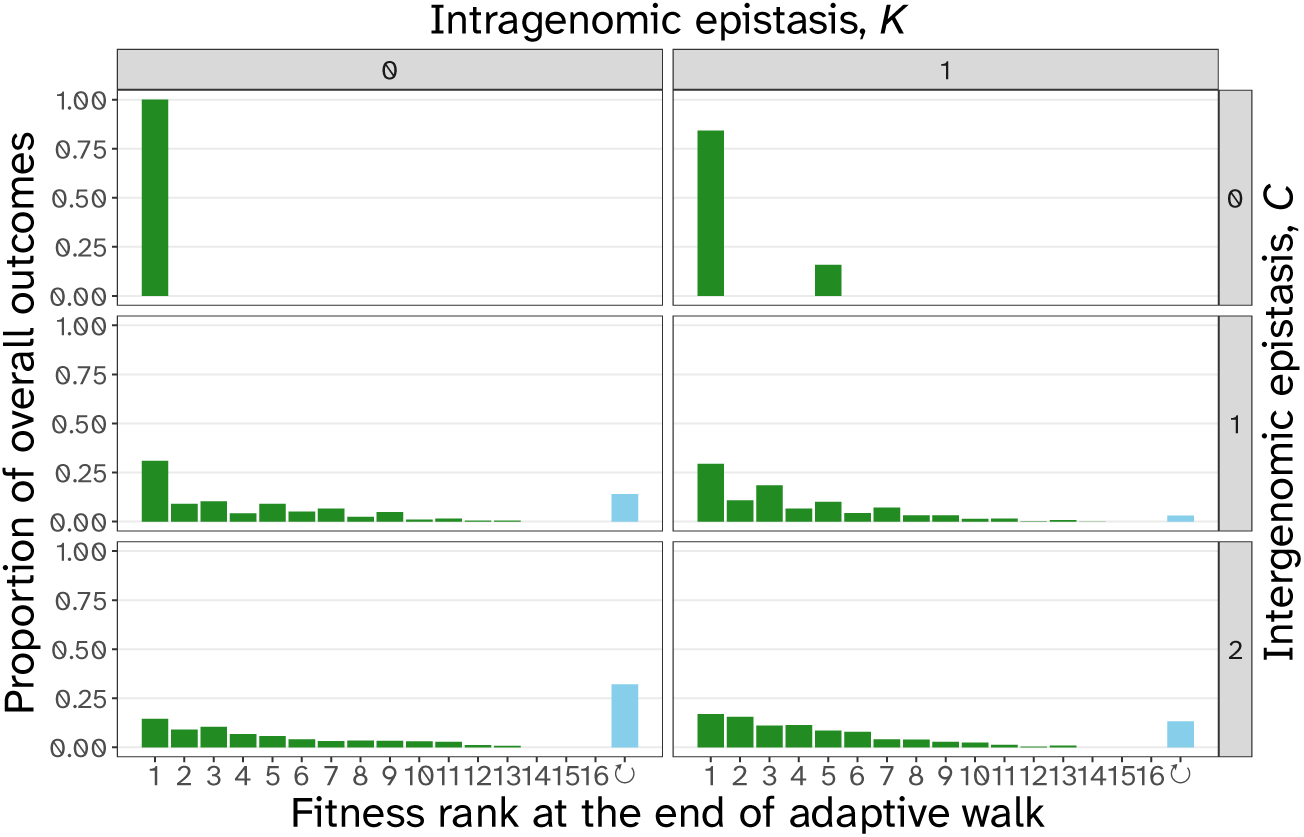
Distribution of observed outcomes of adaptation across all simulated fitness landscapes weighted by their occurrence for *N* = 2. The probability of reaching the global fitness peak (fitness rank 1) is strongly reduced on landscapes with *C ≥* 1 relative to landscapes with no intergenomic epistasis of both *N* = 2 and *N^′^* = 4 (Fig. B.14). The probability of cycling increases with high intergenomic epistasis (*C* = 2), but is reduced by intragenomic epistasis (*K* = 1).

**Figure B.11:**
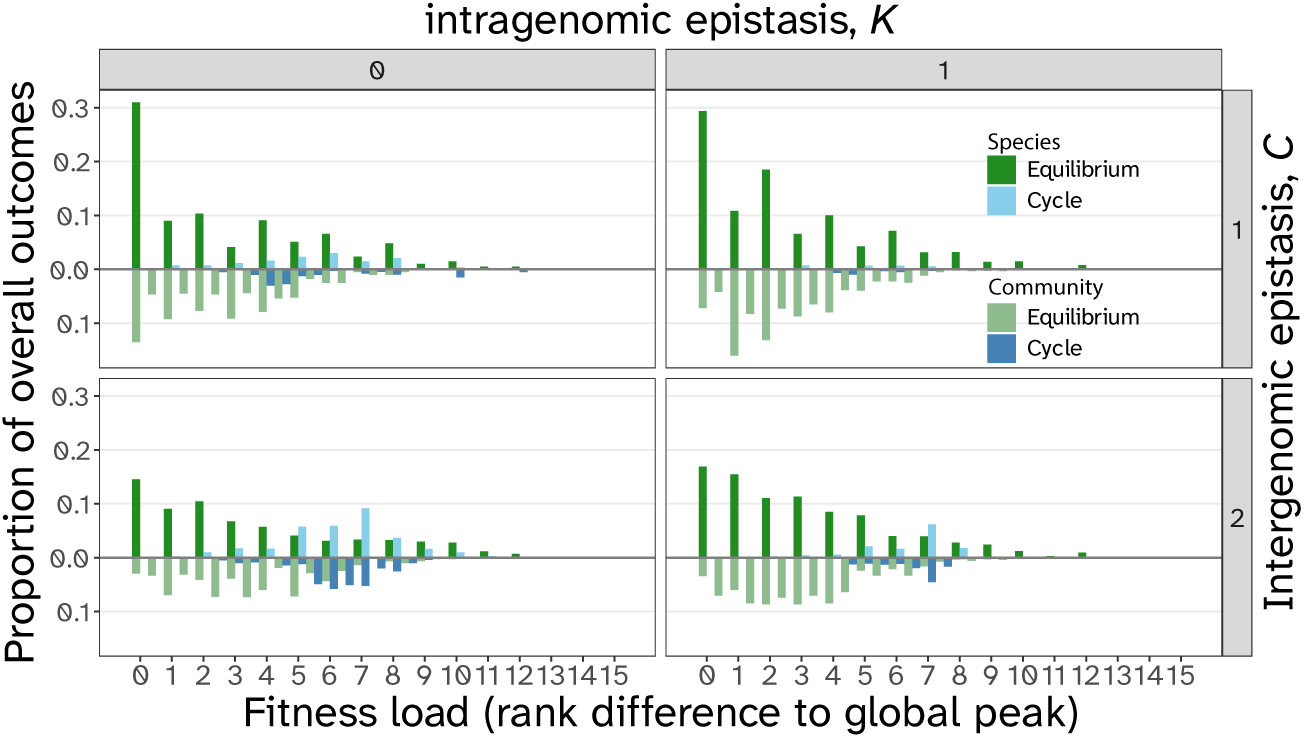
Difference in fitness load between adaptive outcomes at equilibrium (green) and cycles (blue), for individual species (top half of facet) and communities (bottom half of each facet; as quantified by the average fitness load of the interacting species – note that this measure runs in intervals of 0.5 for two-species communities) for *N* = 2. For cycles, we calculate the mean fitness load of each species during the cycle (rounded to the next-higher load category in the histogram). Fitness load is calculated as the rank difference to the global fitness peak (rank 1). The paucity of communities with 0 load arises because interactions are rarely win-win, where both species reach their global fitness peak; instead, there is usually some level of mismatch between the species. Cycles are associated with increased fitness load, with similar distributions of fitness load at the community and at the species levels.

### Single-species comparisons with N^′^ = 4 loci

When comparing *NKC*(*S*) fitness landscapes constructed with *N* = 2, we run into the issue that intro-ducing intergenomic epistasis (C ≠ 0) expands the parameter space and thus comparing landscapes with *C* = 0 to landscapes with C ≠ 0 is limited in certain aspects. Though we believe that this expansion in parameter space by itself is an important outcome to consider when trying to understand the conse-quences of intergenomic epistasis, we also want to provide additional references with a more comparable parameter space. For this, we also simulated single-species fitness landscapes (static environment) and seascapes (dynamic abiotic environment) with *N^′^* = *N* +*N* = 4. See supplementary materials A for more details on the seascapes model we implemented (the *NK*(*E*) model).

**Figure B.12:**
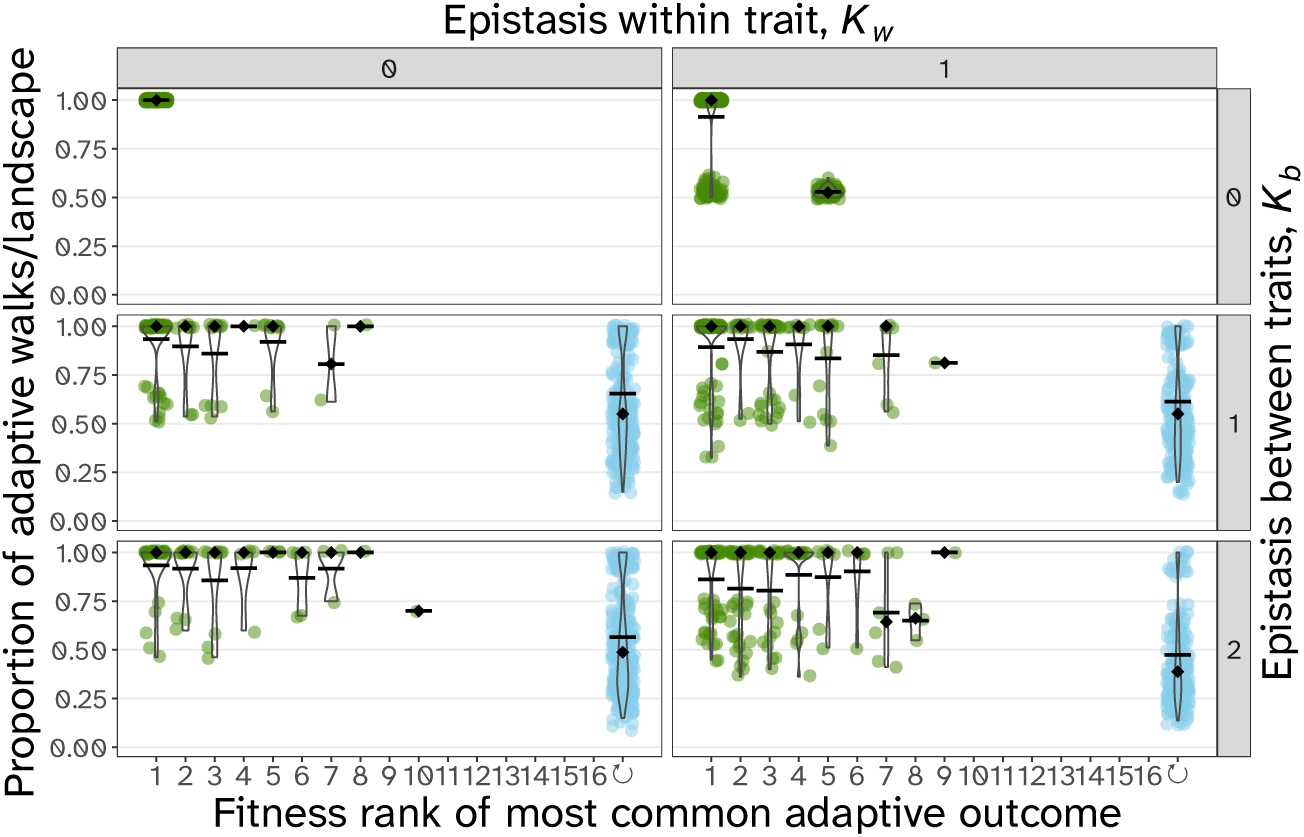
Frequencies of the most common adaptive outcomes in the *NK*(*E*) seascapes model. The most common outcome under the assumption of frequent environment switching is cycling (blue dots; also see Fig. B.13).

**Figure B.13:**
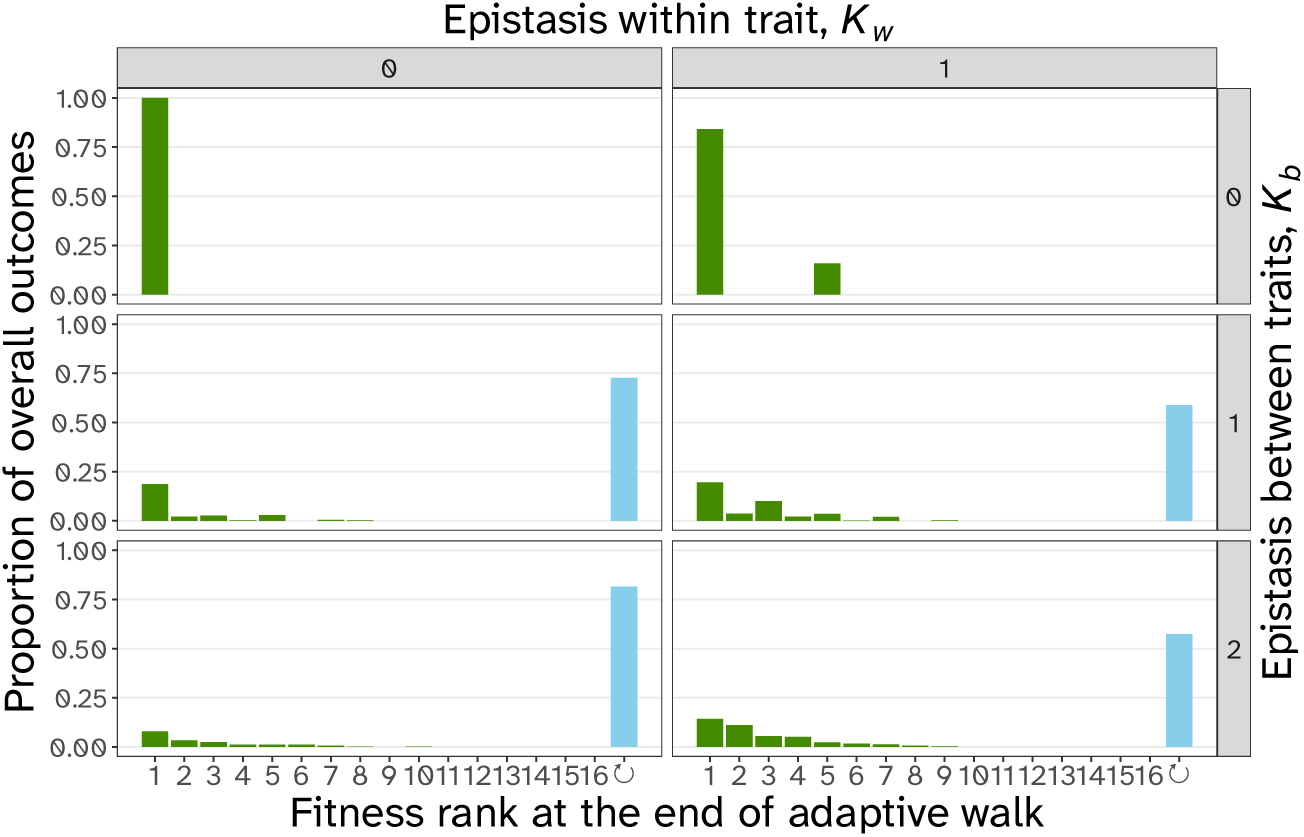
Distribution of observed outcomes of adaptation in the *NK*(*E*) model across all simulated fitness seascapes weighted by their occurrence for *N^′^* = 4 and differing *K_w_* and *K_b_*. The most common outcome of adaptation is cycling between genotypes.

**Figure B.14:**
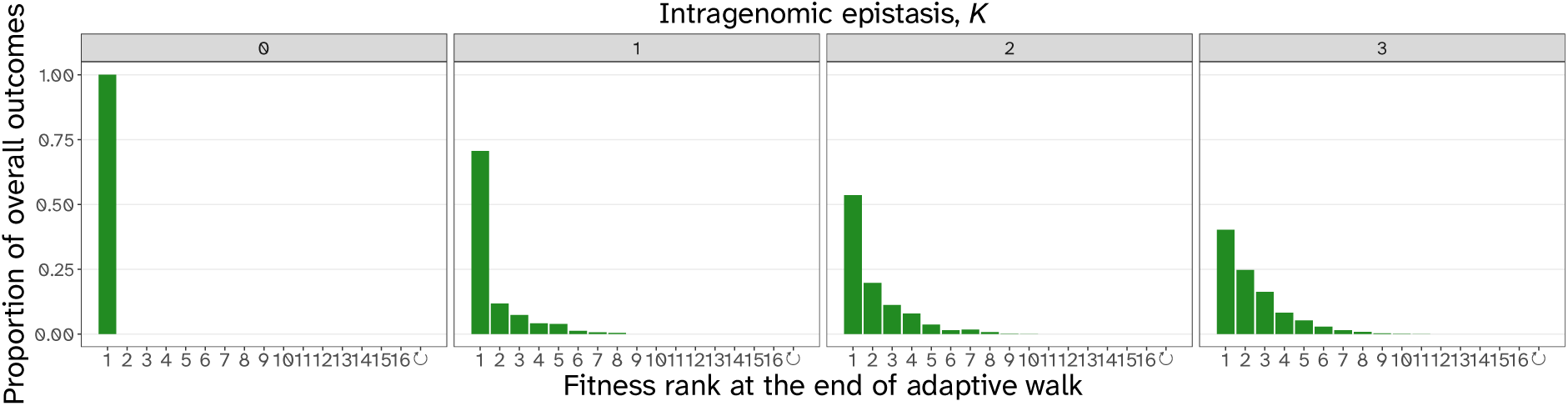
Distribution of observed outcomes of adaptation in the *NK* model across all simulated fitness landscapes weighted by their occurrence for *N^′^* = 4 and *K^′^* = *K* + *C*. The fitness distributions for *N^′^* = 4 match the pattern of repeatability in Fig. 5b, where the global fitness peak is the most common outcome and less fit genotypes are less common.

Similar to the *NKC*(*S*) model we compared different types of epistatic interaction patterns in the *NK* model. The types of interaction matrices we implemented were (1) random interactions - unstructured interactions between the *N^′^* = 4 loci in the genome, (2) modular interactions - interactions are arranged in blocks of *K* + 1 loci, (3) nearest neighbour interactions, (4) preferential interactions - loci that already have many interactions are more likely to add a further interaction than loci that have few interactions (total of *N* × *K* interactions among loci), and (5) hierarchical interactions - few loci influence all other loci (total of *N* × *K* interactions among loci). See Rivkin and Siggelkow (2007) for explanations on preferential and hierarchical interactions. The different interactions matrices affect patterns of endpoint repeatability through the number of observed outcomes on a landscape and the frequency of the most common outcome. This is most notably the case with the hierarchical interaction patterns.

**Figure B.15:**
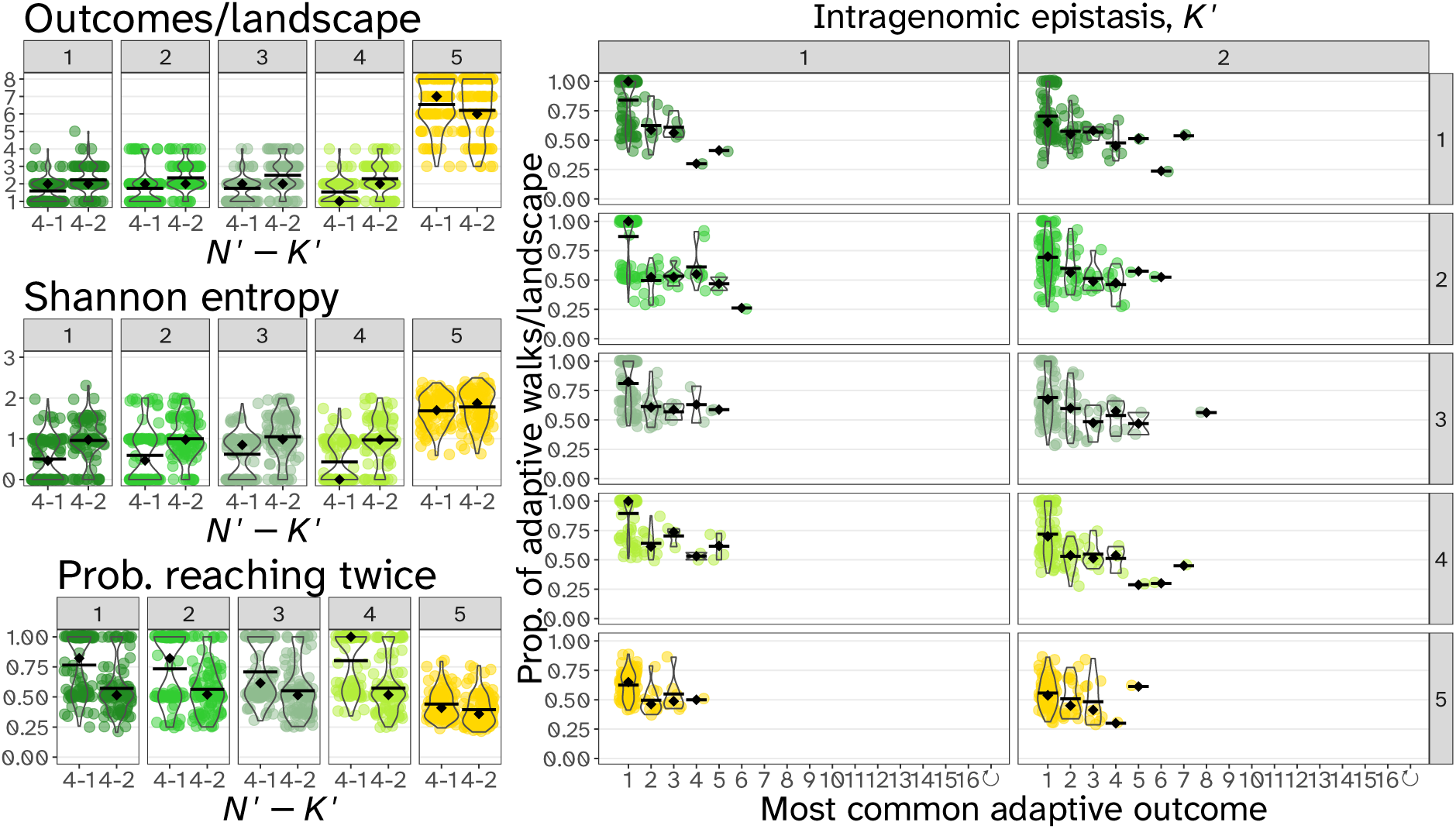
On *NK* landscapes with *K^′^* = 1, 2 we implemented different kinds of interaction matrices. We implemented (1) random interactions - unstructured interactions between the *N^′^* = 4 loci in the genome, (2) modular interactions - interactions are arranged in blocks of *K* + 1 loci, (3) nearest neighbour interactions, (4) preferential interactions - loci that already have many interactions are more likely to add a further interaction than loci that have few interactions (total of *N × K* interactions among loci), and (5) hierarchical interactions - few loci influence all other loci (total of *N × K* interactions among loci). Landscapes generated using genetic interactions following a hierarchical structure (interaction matrix 5) showed decreased endpoint repeatability.

### Differences in mutation rates have little effect on repeatability and variability

**Figure B.16:**
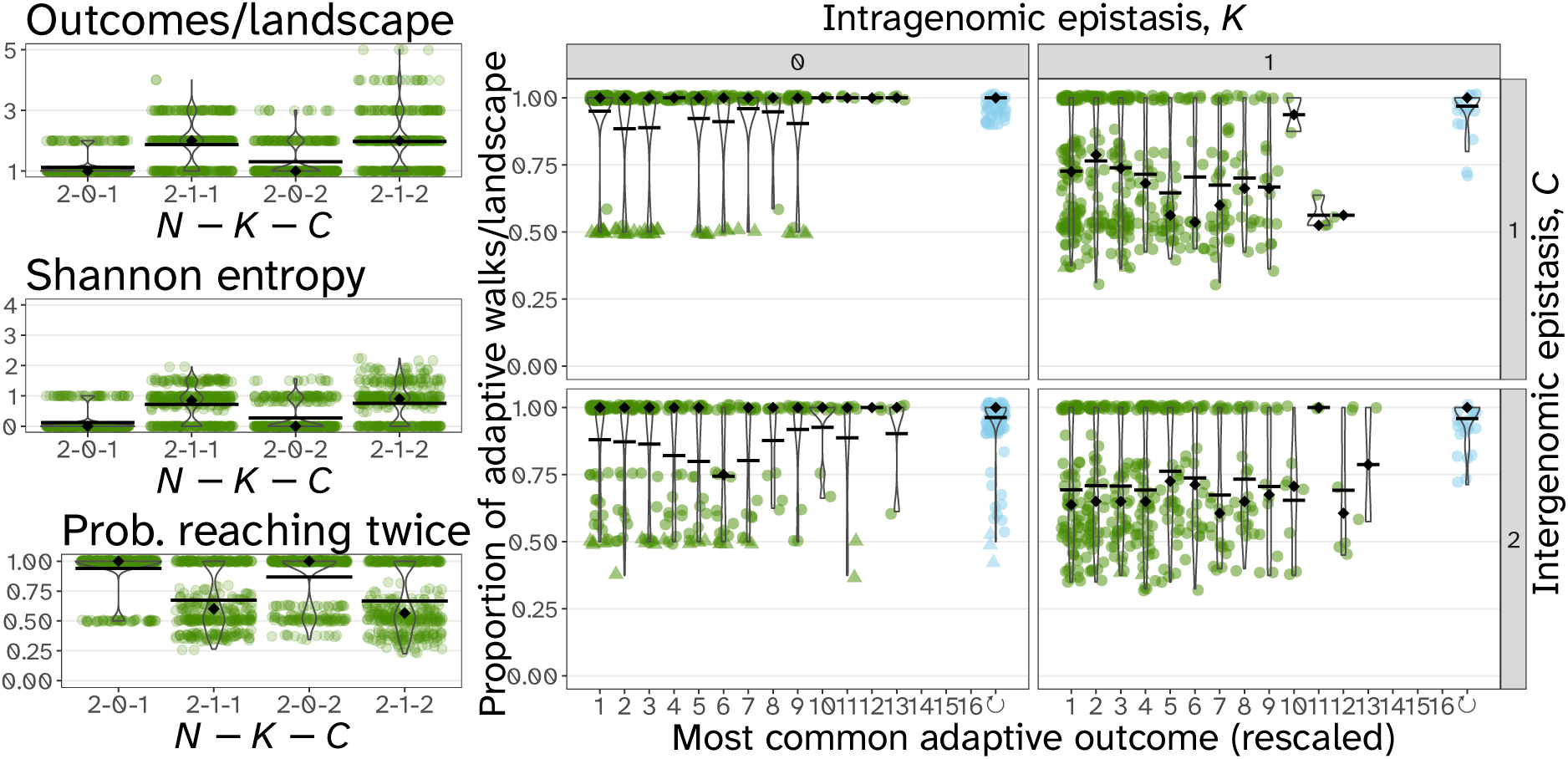
Repeatability results for *N* = 2*, C >* 0 and *R* = 2, where the focal species can take two mutational steps at a time relative to the partner species. Patterns of repeatability are seemingly identical to landscapes with *R* = 1.

**Figure B.17:**
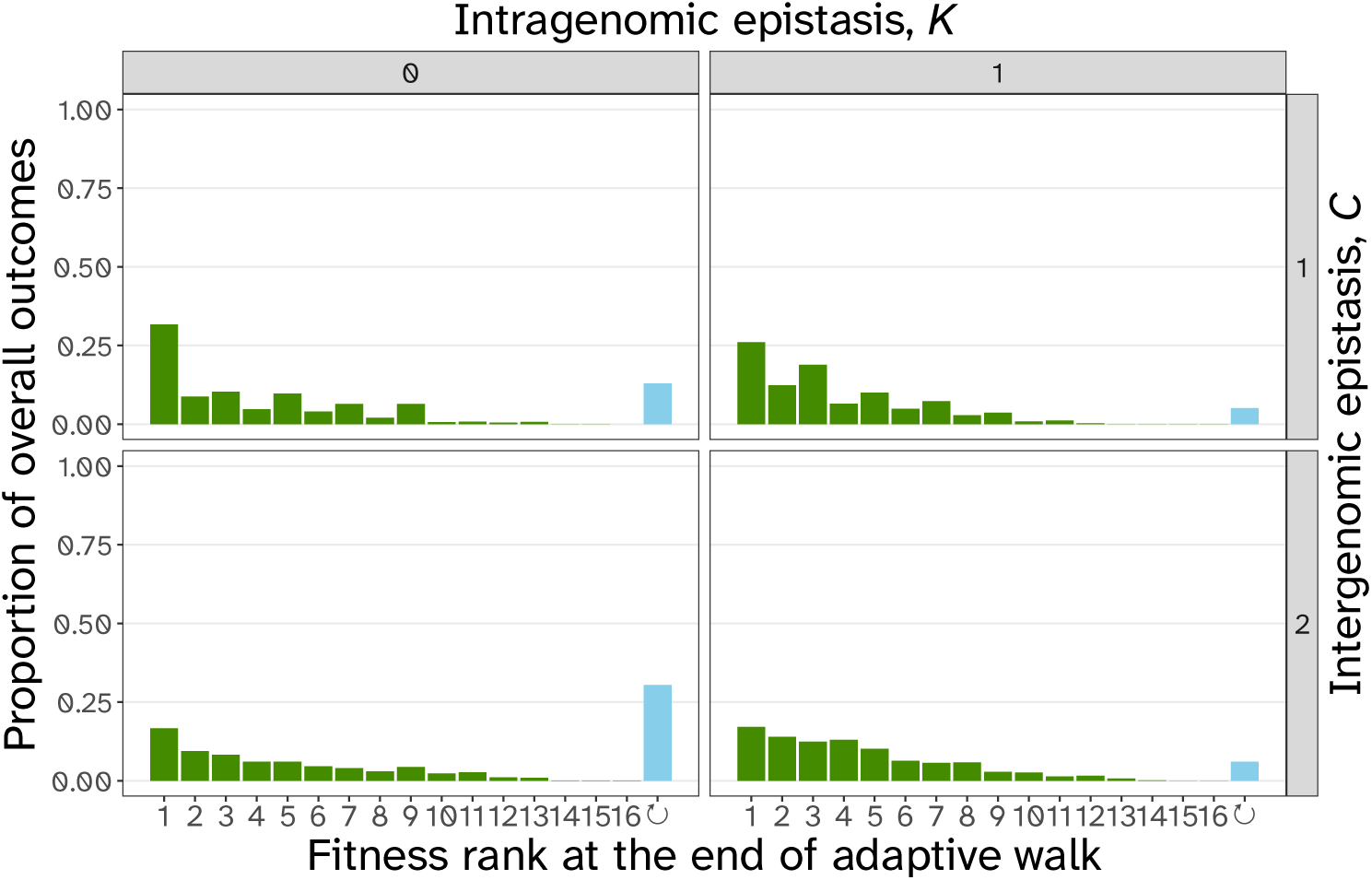
Fitness distribution for *N* = 2*, C >* 0 and *R* = 2, where the focal species can take two mutational steps at a time relative to the partner species. Patterns of fitness distribution are seemingly identical to landscapes with *R* = 1.

**Figure B.18:**
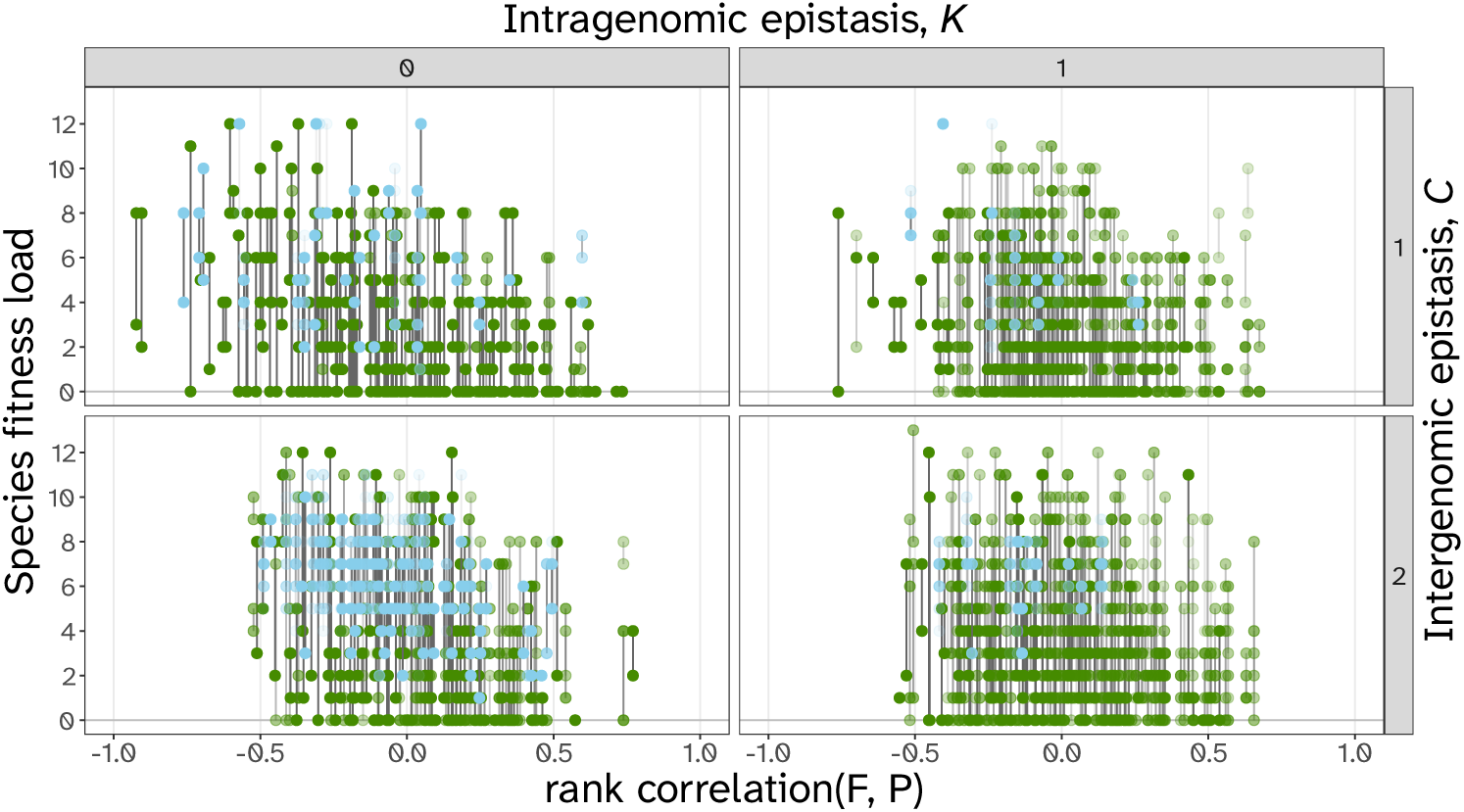
Asymmetry in the mutation rate for *N* = 2*, C >* 0 and *R* = 2 mirrors the results of strictly symmetric coevolution in Fig. B.20 for *R* = 1. Each dot shows the fitness load of a species at the end of an adaptive walk. Species pairs from the same fitness landscape are connected by a vertical grey line. The opacity of the dots corresponds to their level of repeatability.

### The effects of randomness in mutation rate

By introducing a random mutation sequence between both species, the time to equilibrium is substantially increased (*cf.* Kauffman and Johnsen, 1991). Therefore, we are no longer able to distinguish between true cycles and adaptive walks that just have not reached equilibrium yet using our simple method of detecting cycles (cut-off after max. time step). Because of this we have extended the maximum steps in an adaptive walk to 1,000 steps, but then still only report endpoint repeatability for landscapes where all adaptive walks have reached equilibrium (which is 80.875 % of landscapes).

**Figure B.19:**
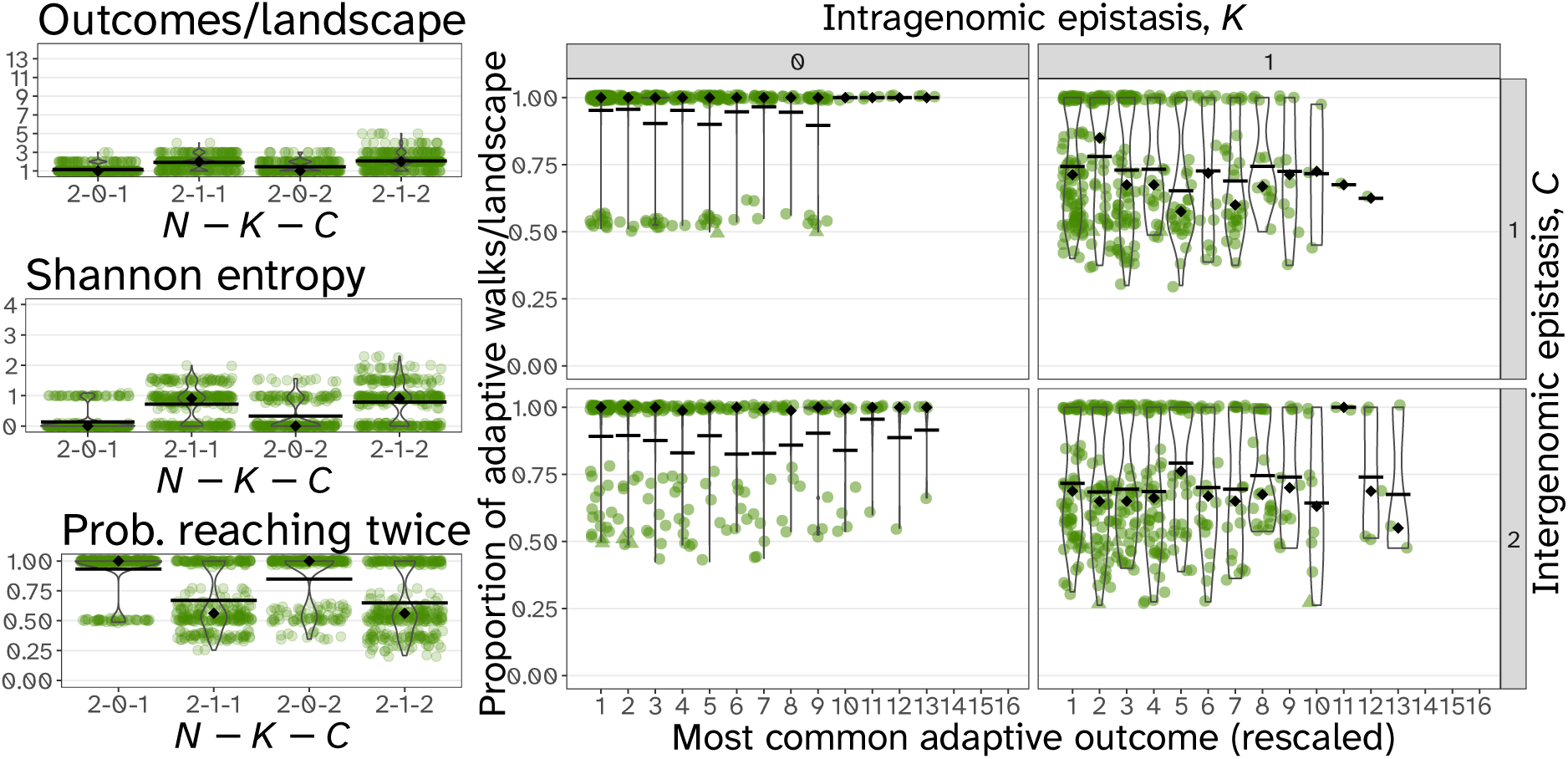
Repeatability results for *N* = 2*, C >* 0 and *R_X_ ∼* Pois(1),. Patterns of repeatability are comparable to landscapes with *R* = 1. Note that we only report statistics based on landscapes that reached stable-state equilibria due to the extended adaptive walks caused by the random mutation pattern.

### Effect of inferred ecological interactions

**Figure B.20:**
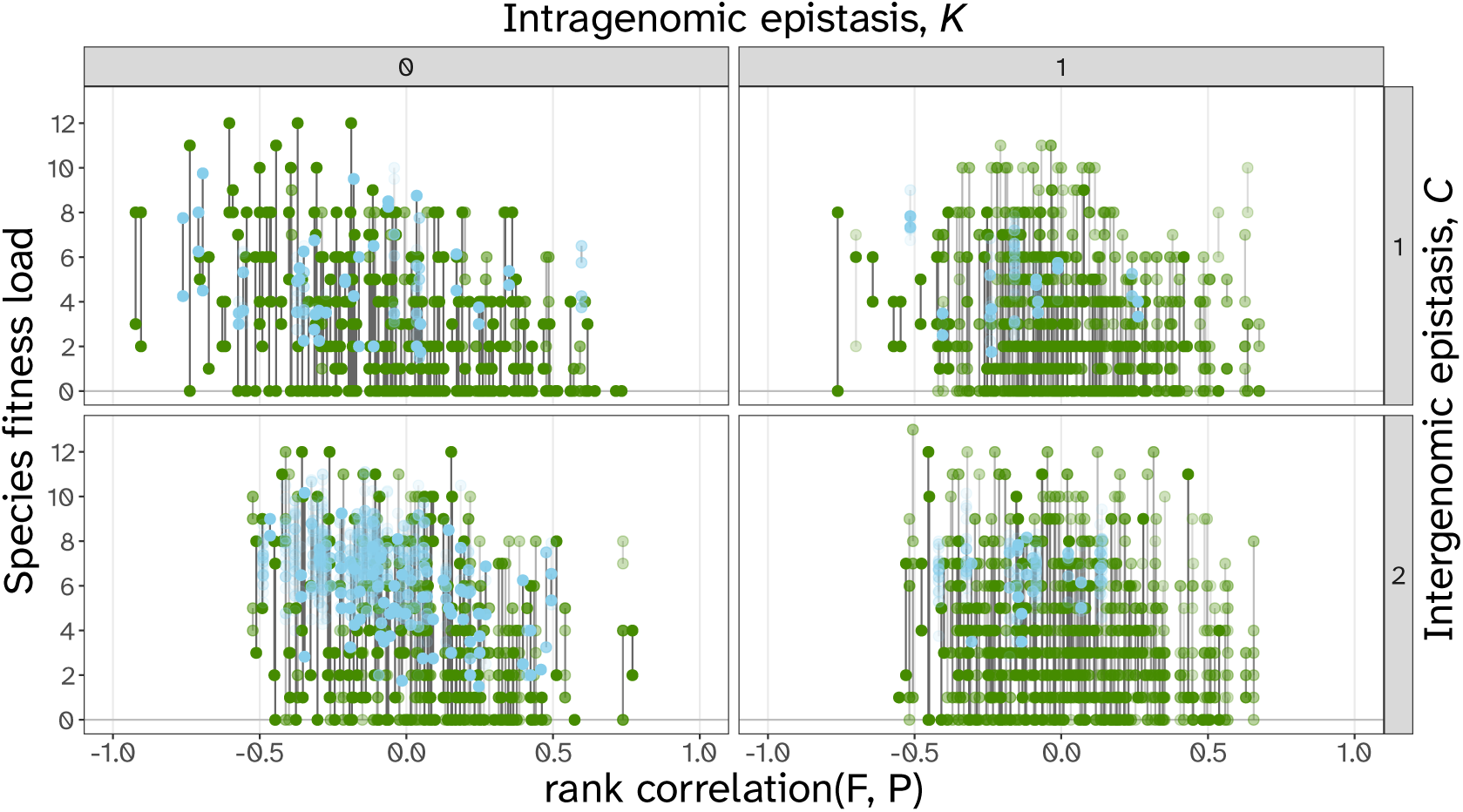
Different results of ecological interactions along the continuum of rank correlations between species with landscapes of size *N* = 2. Each dot shows the fitness load of a species at the end of an adaptive walk. Species pairs from the same fitness landscape are connected by a vertical grey line. The opacity of the dots corresponds to their level of repeatability. Green shows equilibrium results at joint fitness peaks, and blue shows the mean fitness of species during cycles.

In order to understand the trends in fitness load associated with coevolution, we fit two generalised linear mixed effect models for the differences in fitness rank between species and the community fitness load (mean fitness load between species). The model outputs generated with the *R* package *glmmTMB* (version 1.1.9; Brooks et al., 2017) can be found in Table B.1.

**Table B.1:** We fit generalised linear mixed effect models using the *R*package *glmmTMB* (version 1.1.9; Brooks et al., 2017) to estimate how the difference between fitness ranks and the community fitness load change across the rank correlations between species. The formula we use is “response *∼* rank correlation + c + k + cycle” where we weight the outcomes by how often they occur.

| Response |  | Estimate | SE | Z | p |
| --- | --- | --- | --- | --- | --- |
| *Difference in fitness rank* | Intercept | 0.820555 | 0.008072 | 101.66 | $< 2e - 16$ |
| | Rank correlation | -0.872592 | 0.008278 | -105.42 | $< 2e - 16$ |
| | $C$ | 0.191821 | 0.004859 | 39.47 | $< 2e - 16$ |
| | $K$ | -0.110231 | 0.004868 | -22.64 | $< 2e - 16$ |
| | cycle-yes | -0.606267 | 0.008031 | -75.49 | $< 2e - 16$ |
| *Community fitness load* | Intercept | 0.870693 | 0.006871 | 126.72 | $< 2e - 16$ |
| | Rank correlation | -0.595772 | 0.006928 | -85.99 | $< 2e - 16$ |
| | $C$ | 0.288781 | 0.004056 | 71.20 | $< 2e - 16$ |
| | $K$ | -0.020283 | 0.004029 | -5.03 | $4.81e - 07$ |
| | cycle-yes | 0.476656 | 0.004816 | 98.98 | $< 2e - 16$ |

**Figure B.21:**
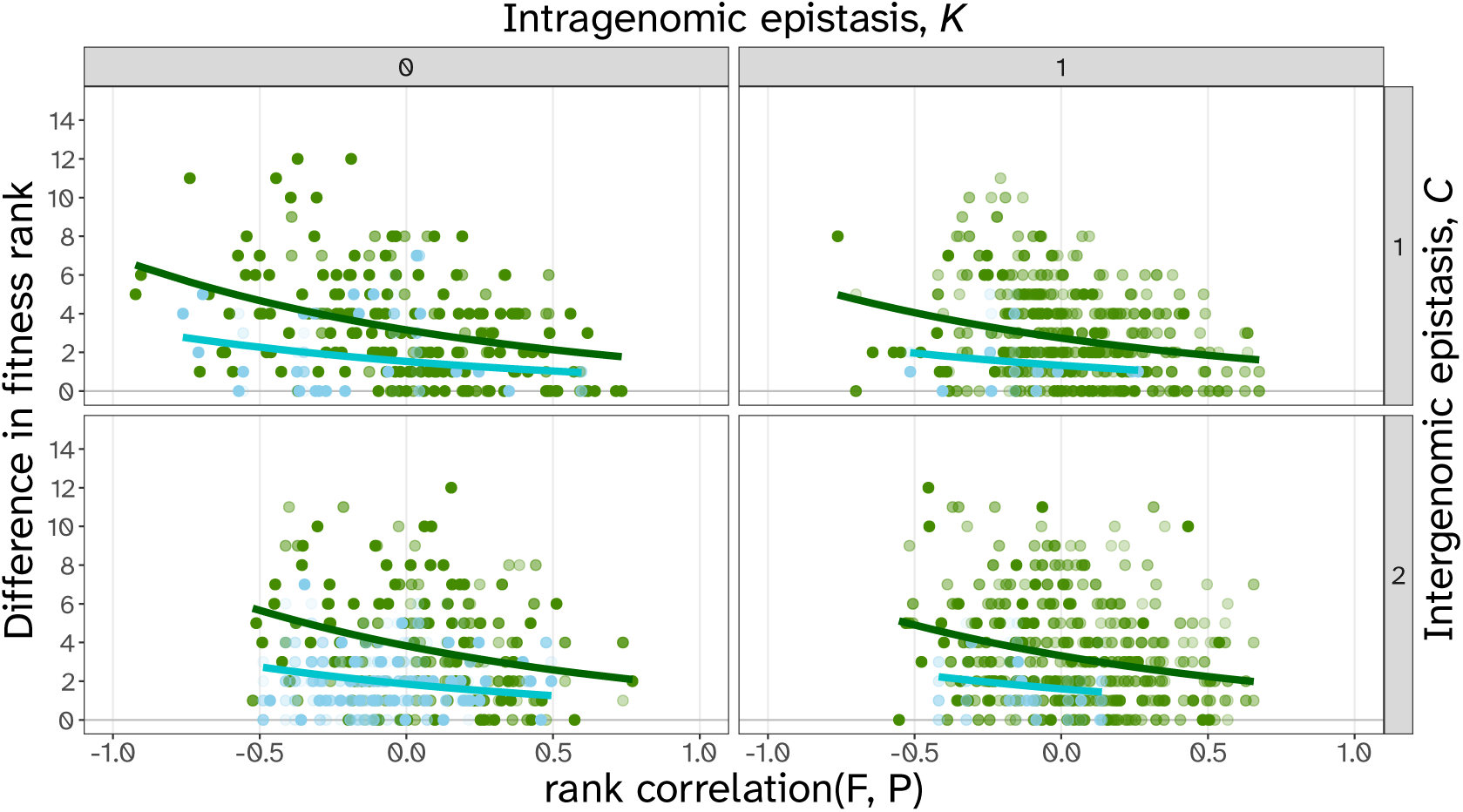
Difference in fitness rank between both species across the rank correlations between species. Each dot shows the difference in fitness load of both species at the end of an adaptive walk. The opacity of the dots corresponds to their level of repeatability. Green shows equilibrium results at joint fitness peaks, and blue shows the cycle results based on mean fitness during the cycle. See Table B.1 for fit of generalised linear mixed model.

**Figure B.22:**
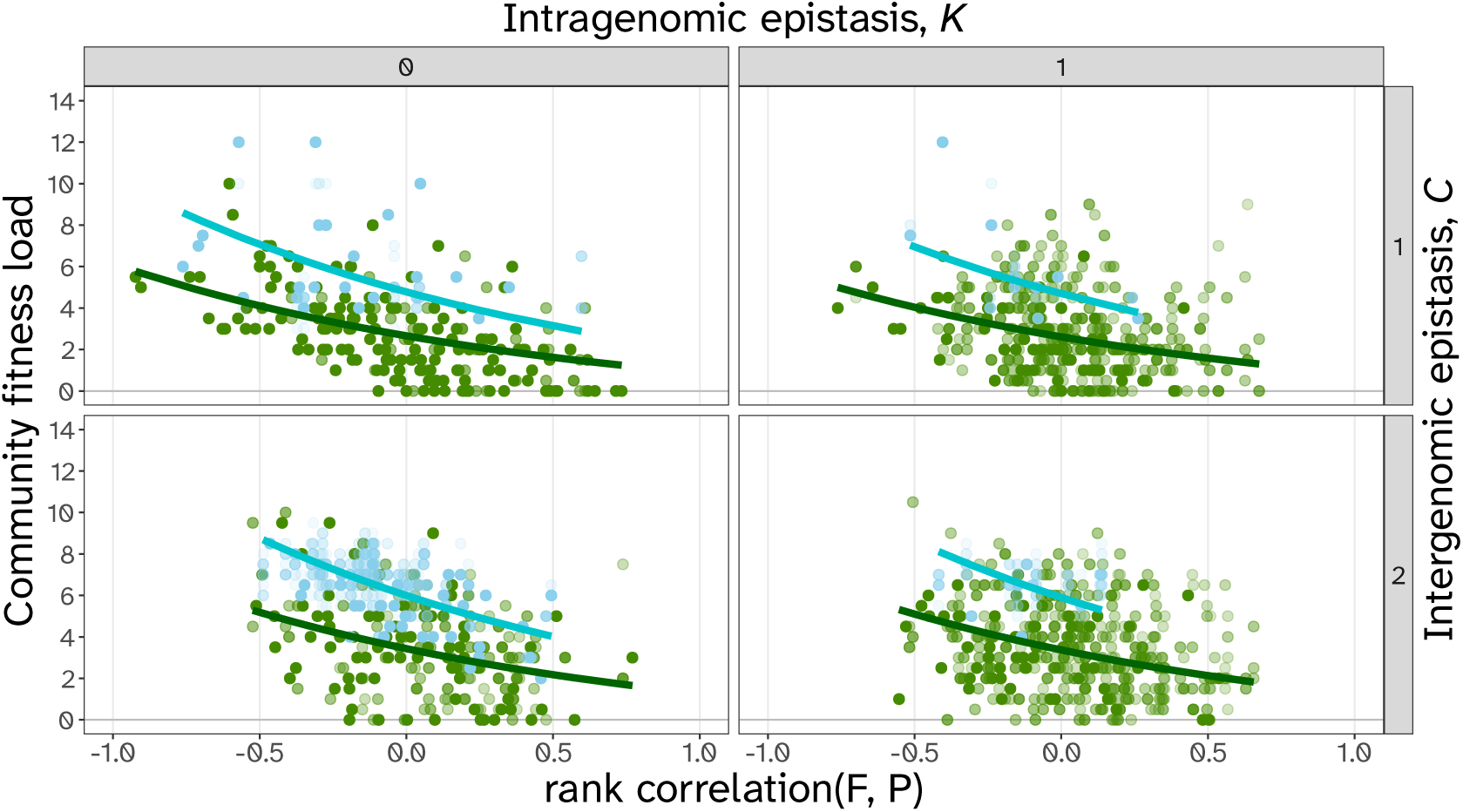
Community fitness load across the rank correlations between species. Each dot shows the mean community fitness rank at the end of an adaptive walk. The opacity of the dots corresponds to their level of repeatability. Green shows equilibrium results at joint fitness peaks, and blue shows the cycle results based on mean fitness of each species during the cycle. See Table B.1 for fit of generalised linear mixed model.

### Inferred landscape properties do not correlate with endpoint repeatability

**Figure B.23:**
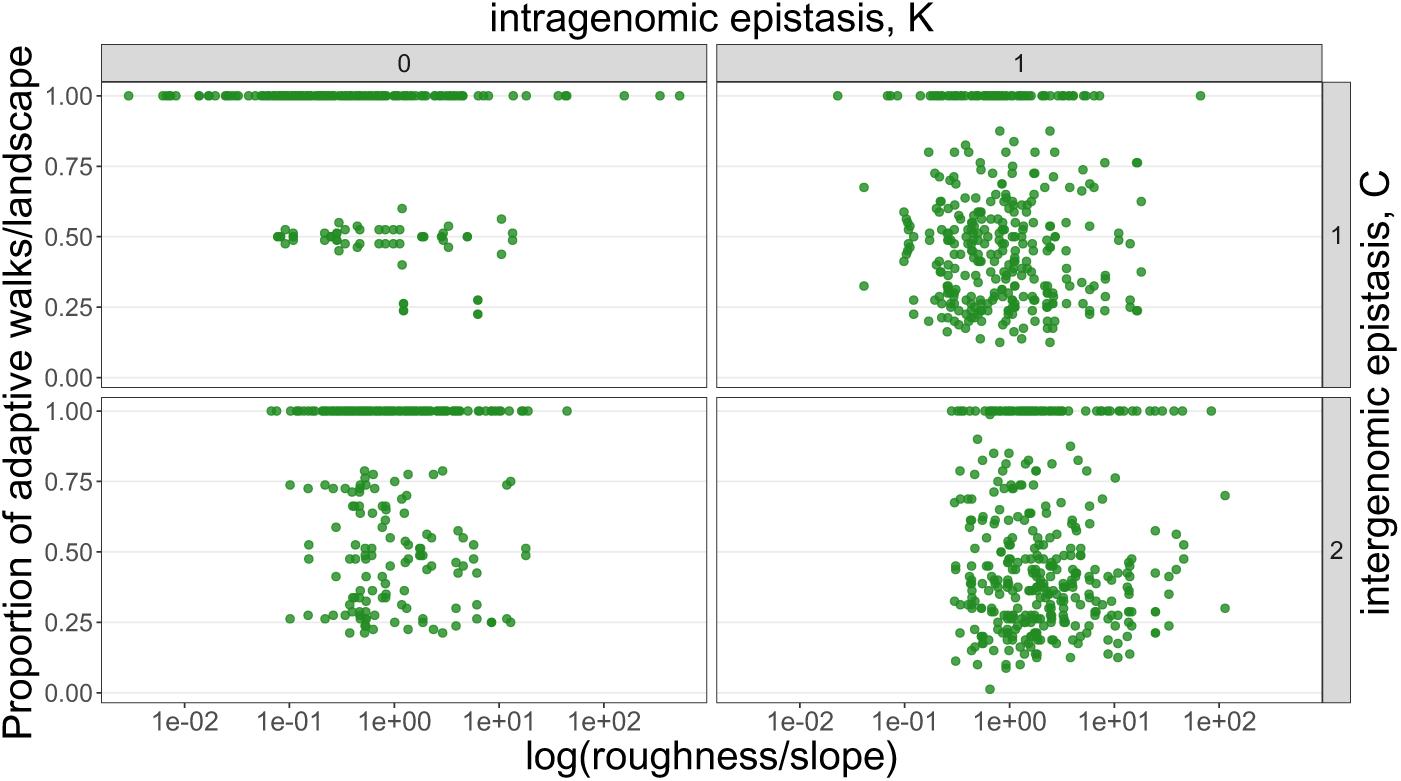
Endpoint repeatability of different adaptive outcomes for *N* = 2*, C >* 0 is not explained by the inferred ruggedness (roughness to slope ratio) of the fitness landscape.

**Figure B.24:**
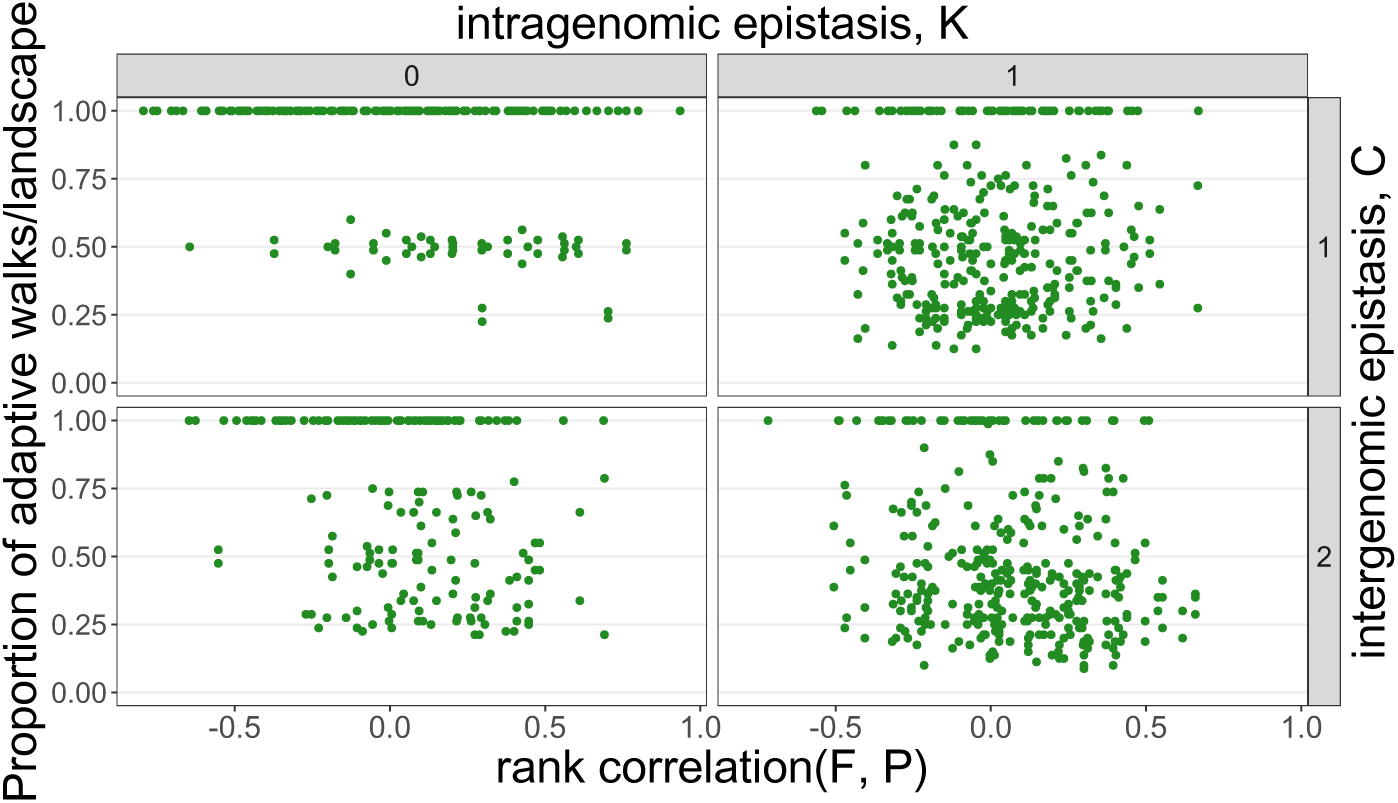
Endpoint repeatability of different adaptive outcomes for *N* = 2*, C >* 0 does not depend on the correlation of fitness ranks between species.

**Figure B.25:**
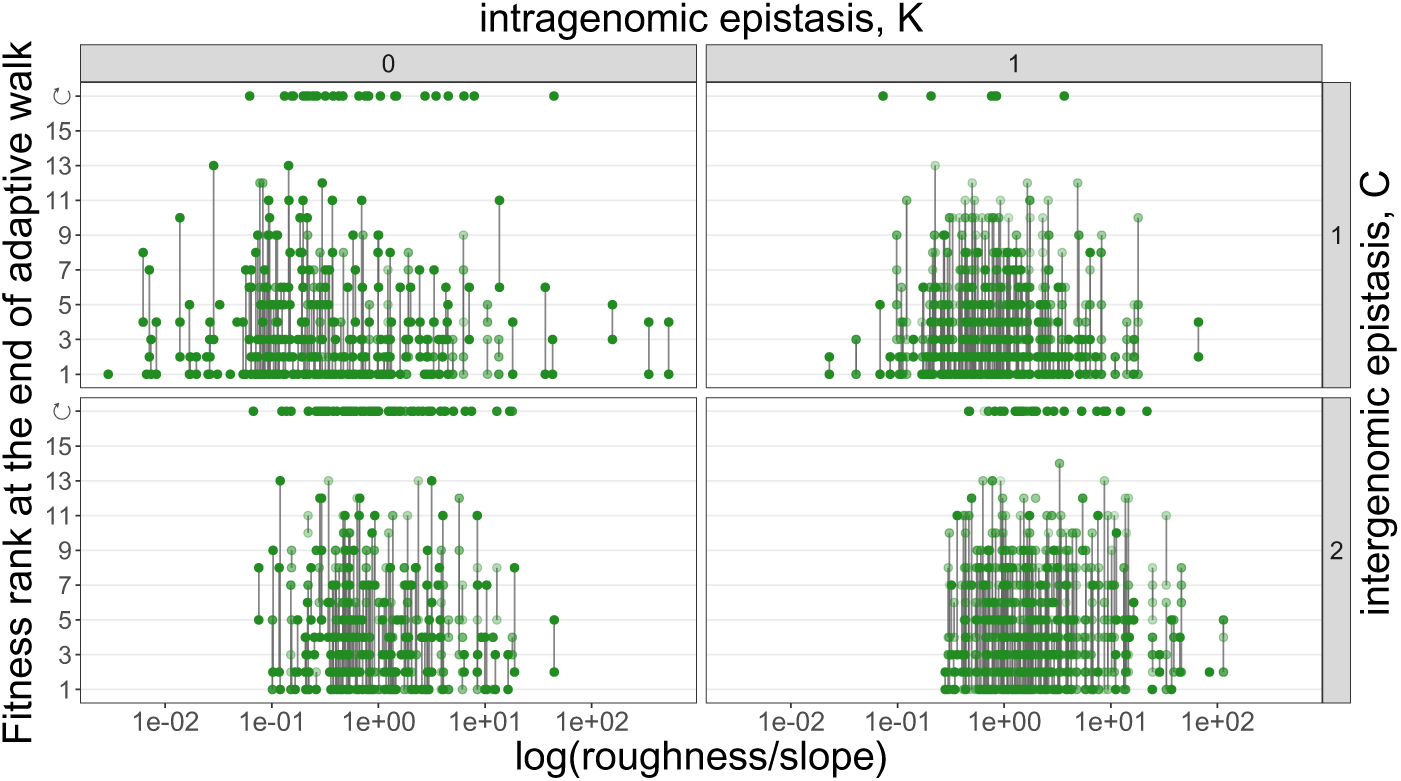
Interaction results and cycling for *N* = 2*, C >* 0 do not depend on the inferred ruggedness (roughness to slope ratio) of the fitness landscape.

